# Netrin-1 drives cell-type-specific plasticity of human dopaminergic neurons during circuit integration in a Parkinsonian model

**DOI:** 10.64898/2026.06.01.729439

**Authors:** Chiara Pavan, Kristian Honnens de Lichtenberg, Carlos W Gantner, Stefano Frausin, Joseph Chen, Cameron PJ Hunt, Brianna Xuereb, Emily Hart, Zac Chatterton, Glenda Halliday, Niamh Moriarty, Clare L Parish

**Author notes:** Correspondence: Chiara Pavan, Clare Parish (Lead author).

## Abstract

The survival of transplanted ventral midbrain (vm) dopaminergic neurons (DAn) and their innervation of host striatal tissue are crucial for ameliorating motor symptoms in Parkinson’s disease (PD). However, human pluripotent stem cells (hPSC) show inferior axonal plasticity compared to fetal donor tissue. While modulation of the host environment with trophic cues, such as glial cell-derived neurotrophic factor (GDNF), can improve graft outcomes, these cues lack specificity for DAn, resulting in plasticity of other neurons within the graft. Using single nuclei RNA sequencing, we identified axonal guidance pathways (Semaphorin, Netrin and Wnt) that were preferentially activated in DAn within the graft. Overexpression of Semaphorin3A/SEMA3A, Netrin1/NTN1 or WNT5A in the striatum of Parkinsonian mice following vm DA progenitor transplantation promoted A9-DA specification and selectively increased DA innervation of the host striatum, without off-target extrastriatal DA innervation observed in response to GDNF. In Parkinsonian rats, NTN1 overexpression promoted graft-induced motor recovery, selective DA plasticity and activation of postsynaptic striatal neurons without evidence of non-DAn plasticity. Further, snRNA-sequencing of NTN1 or GDNF-treated grafts confirmed the upregulation of DA-specific plasticity by NTN1, while GDNF promoted plasticity in both DA and non-DAn. These findings highlight the capacity to improve on-target integration of hPSC-derived DAn in grafts by selectively targeting DA-specific plasticity. Taken together, these results demonstrate the utility of DA-specific cues to promote functional recovery, improve graft predictability, and limit off-target innervation.

## INTRODUCTION

A prerequisite for neural grafts to ameliorate Parkinsonian motor symptoms is the capacity of the new ventral midbrain dopaminergic (vmDA) neurons to synapse with host striatal neurons ^1–3^. However, following transplantation of vmDA progenitors, the level of reinnervation of the host tissue remains well below that of the intact brain, likely due to the lack of trophic cues in the adult brain, with the noted observation that human pluripotent stem cell (hPSC)-derived vmDA transplants show significantly inferior plasticity compared to fetal tissue grafts ^4^. These observations highlight the need to improve the host environment to encourage plasticity and functional outcomes in graft recipients.

During development, the controlled spatial and temporal release of multiple neurotrophic and morphogenic cues drives neurogenesis, differentiation, migration, axonal pathfinding and survival ^5,6^. Downregulation of these cues in the adult brain results in a less permissive environment for the survival and integration of transplanted neural progenitors ^7,8^. To counteract this, many groups have explored the delivery of developmentally relevant neurotrophic cues. Among these, glial-derived neurotrophic factor (GDNF) has received the greatest attention and has been shown to promote survival, plasticity and metabolism of DA neurons both *in vitro* and *in vivo* (see review ^9^). In the context of neural grafting, GDNF delivery into the host tissue has been shown to enhanced survival, plasticity, and functional integration of DA neurons derived from fetal donor tissue ^10–17^. Similarly, GDNF overexpression in the host striatum, via adeno-associated virus or biomaterial-based delivery, has been shown to influence survival and plasticity of hPSC-derived vmDA neurons ^18–20^. However, GDNF lacks specificity for vmDA neurons, with evidence of GDNF also impacting the survival and plasticity of other neurons within neural grafts ^19^. Thus, alternate trophic cues that selectivity act on vmDA neurons may be superior in promoting striatal targeted functional recovery while minimising off-target effects.

Unfortunately, our knowledge of the regulators of human DA axon guidance has been limited, in part due to the constraints associated with studying human fetal tissue and thereby reliance on other species. In this regard, hPSCs present a valuable tool for advancing our understanding of neural development and can be aided through transplantation whereby new cells can form appropriate synaptic connections not achievable *in vitro*. Transcriptional profiling approaches now enable characterisation of maturing neurons during establishment of defined neural circuits. Whilst we and others have previously profiled hPSC-derived vm progenitor grafts, studies have been limited by: (i) bulk, rather than single cell transcriptional analysis, (ii) under-represented neuronal populations due to harsh mechanical dissociation and/or (iii) lacked single cell resolution ^21–25^. To preserve single-cell level analysis and ensure non-bias profiling within the graft, here we adopted single nuclear RNA-sequencing (snRNA-seq) to profile hPSC-derived vm DA grafts at 12 weeks after transplantation, a period corresponding to active plasticity of the grafted neurons ^21^, screening specifically for axon guidance receptors and/or activated pathways in DAn. We identified 3 pathways - Netrin, Wnt and Semaphorin and subsequently assessed the impact of over-expression of the 3 axonal guidance cues NTN1, WNT5A and SEMA3A on graft maturation and plasticity for both DA and non-DA neurons. We further explored the selective specificity of NTN1 on graft-derived DA and non-DA neurons by single nuclei sequencing and benchmarked this to GDNF. With several hPSC clinical trials for Parkinson’s disease ongoing, our findings highlight significant scope for further refinement through combined cell and gene therapies that selectively promote DA plasticity and recovery of motor function.

## RESULTS

### snRNA-seq identifies xenografted DA neurons and associated genes with potential to selectively modulate their plasticity

The human iPSC line RM3.5-PITX3-GFP was differentiated into VM progenitors, with efficiency of the differentiation confirmed by the co-expression of OTX2, FOXA2 and NESTIN at 13 days in vitro (DIV) and GFP (PITX3), FOXA2 and TH at 25 DIV (Figure 1A). To characterize the transcriptome of transplanted neurons, we grafted vmDA progenitors into rats 2 weeks after unilateral ablation of the nigrostriatal pathway. At 12 weeks, corresponding to a period of active axonal plasticity of the grafts, animals were culled and the striatum, containing the graft, snap frozen. Nuclei were extracted from the frozen tissue and fluorescence-activated nuclei sorting (FANS) used to enriched for single nuclei (Supplementary Figure 1B-D) that were transcriptionally profiled using the 10x platform (Figure 1B). An additional brain verified the presence of a viable graft within the host striatum (haematoxylin and eosin staining, Supplementary Figure 1A).

**Figure 1.**
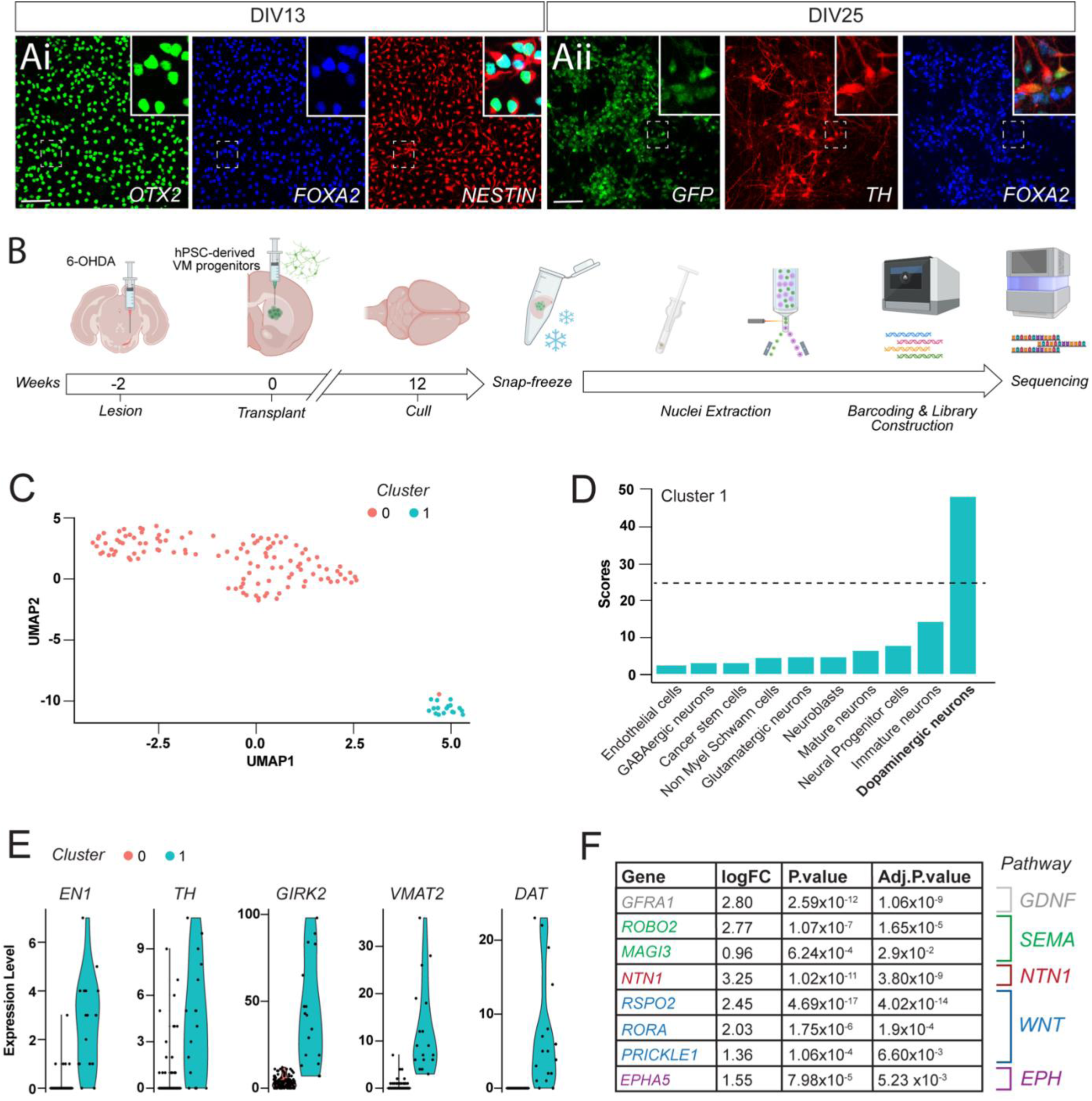
Ventral midbrain dopamine neurons grafts are profiled using single nuclei RNA-sequencing (snRNA-seq). (Ai) Photomicrographs showing high OTX2, Nestin and FOXA2 co-expression within progenitors at 13 days in vitro (DIV) and (Aii) PITX3-GFP, TH and FOXA2 at 25DIV, indicative of VM DA neurons. (B) Schematic summary of in vivo study design for snRNA-seq: 6-OHDA-induced ablation of host vmDA neurons was followed by hPSC-derived vmDA transplantation. Animals were culled at 12 weeks, striatal tissue containing grafts snap frozen, nuclei extracted and stained, prior to snRNA-seq. (C) UMAP embedding with louvain clustering of 133 human single-nuclei identified two distinct clusters. (D) ScType score unambiguously identified cluster 1 as DA neurons. Dotted line indicates thresholds for cell assignment. (E) Violin plot showed higher expression of DA-associated genes in cluster 1 than in cluster 0. (F) logFC, P-value and adjusted P value of upregulated genes within the xenograft DA that belonged to GDNF, semaphorin (SEMA), WNT, netrin (NTN1) and ephrin (EPH) signalling pathways. Scale bars: (A) 100 μm. Abbreviations: 6-OHDA, 6-hydroxydopamine; DIV, days in vitro; FANS, fluorescence activated nuclei sorting; GFP, green fluorescent protein; GDNF, glial derived neurotrophic factor; GO, gene ontology; H&E, hematoxylin and eosin; hiPSC, human induced pluripotent stem cells; HNA, human nuclear antigen; NTN, netrin; NeuN, Neuronal Nuclei; SEMA, semaphorin; Tx, transplantation.

Sequenced samples were aligned to a hybrid rat and human genome and nuclei in which >90% of mRNA were aligned to human transcripts were denoted as human origin. These human nuclei represented a small percentage (4.2% and 2.1% of the total nuclei for sample 1 and sample 2 respectively), amounting to a total of 133 nuclei (Supplementary Figure 2C). As only minor batch effect was found (Supplementary Figure 2D), the two samples were pooled. Re-clustering of the human nuclei identified two distinct clusters, cluster 0 and cluster 1, with 116 and 17 cells respectively (Figure 1C). Neither %mt RNA (Supplementary Figure 2E) or number of genes detected per cell (Counts, Supplementary Figure 2F) differed between the two clusters. Using cell-type reference gene sets (scType), the identity of cluster 1 was readily ascribed as DA neurons (Figure 1D), while cluster 0 comprised a heterogeneous population (Supplementary Figure 2G).

To identify potential neurotrophic targets specific to DA neurons, we performed differential expression analysis between DA (Cluster 1) and non-DA nuclei (Cluster 0). We identified 502 differentially expressed genes (DEG’s; adj. p-value <0.05) of which 202 were upregulated and were enriched for gene ontologies related to neuronal structure and function (Supplementary Figure 2H). As expected, vmDA markers (EN1, KCNJ6/GIRK2 and TH) were upregulated in the graft-derived DA neurons as well as neurotransmitter receptors and transporters (DRD2, SLC18A2/VMAT2 and SLC6A3/DAT) (Figure 1E). Upregulated genes within graft-derived DA neurons were significantly enriched for genes predicted to be a part of the “Synaptic Membrane” (adj-val = 0.00009, Supplementary Figure 2H). Further screening for genes associated with axon guidance and plasticity revealed the following differentially expressed genes of interest: GFRα1 (a GDNF receptor); ROBO2 and MAGI3, involved in axon guidance receptor activity linked to the semaphorin (SEMA) pathway; RSPO2, RORA and PRICKLE involved in WNT signalling; NTN1, implicated in netrin (NTN) signalling; and finally EPHA5, linked to EPHRIN signalling, Figure 1F. These pathways (SEMA, NTN, WNT and EPHRIN) can be initiated by (but not limited to) SEMA3A, NTN1, WNT5A and EPHRINA5, respectively, with multiple studies showing these axonal guidance molecules support neurite extension and/or guidance during the development of the nigrostriatal pathway ^26–35^. Noting challenges in sustaining Ephrin over-expression to guide axon pathfinding and plasticity *in vivo*, due to their membrane bound ligand and receptor, here we focused our attention on the 3 key secreted guidance cues, SEMA3A, NTN1 and WNT5A, to assess their ability to more selectively regulate plasticity of DA neurons within hPSC-derived neural grafts, in comparison to GDNF.

### Exposure of vm progenitor grafts to SEMA3A, NTN1 or WNT5A improves DA maturation

To achieve targeted over-expression of the identified guidance cues, we engineered adeno-associated viral (AAV) vectors carrying the full-length coding sequence (CDS) of either SEMA3A, NTN1 or WNT5A, as well as an empty vector and GDNF for comparative purposes. Unlike GDNF, for which reliable antibodies exist, viruses containing the NTN1, SEMA3A or an empty vector included a fluorescence reporter (mCherry) to enable immunohistochemical validation of expression in the host brain, Supplementary Figure 3A-D. This was not feasible for SEMA3A due to the large size of the gene, noting inclusion of the reporter could negatively impact viral transduction efficiency. Transduction efficiency of the viral vectors was therefore additionally verified by real-time PCR of homogenized tissue from mice that received striatal AAVs injections (Supplementary Figure 3E).

To assess the impact of SEMA3A, NTN1 and WNT5A on graft plasticity, 6-OHDA lesioned mice were transplanted with PITX3-GFP hiPSC-derived VM progenitors and 2 weeks later viral vectors delivering the axonal guidance proteins were injected into the host dorsolateral striatum (Figure 2A). At 24 weeks post-transplantation, all animals had surviving grafts, identified as dense hNCAM immunoreactive deposits within the striatum (Figure 2B-F), and were confirmed to have sustained expression of NTN1, WNT5A or GDNF in the dorsolateral striatum (by mCherry or GDNF immunohistochemistry, data not shown). No aberrant neural overgrowths/tumour-like formations were observed. Volumetric assessment showed that exposure to GDNF significantly increased graft size, with trending volume increased observed for SEMA3A and WNT5A, but not NTN1 exposed grafts (Figure 2G). Unsurprising, given the delayed delivery of the viral vectors/protein expression relative to timing of cell implantation, the total number of human cells (HNA^+^, Figure 2H, K), human neurons (HNA^+^ NEUN^+^, Figure 2I, K) and the proportion of human neurons (% NEUN^+^HNA^+^/HNA^+^, Figure 2J, K) were not different between groups. GFP^+^ staining revealed the clustering of vmDA neurons (GFP^+^) at the border of the transplants (Figure 2L-P,L’), as previously observed in fetal and PSC-derived transplants ^3,18^. Despite differences in graft volume, the total number of GFP^+^ cells was similar across the groups, suggestive that GDNF influenced the dispersion of cells within the graft (Figure 2Q-R). We next assessed the impact of trophic cue overexpression on DA subtype identity. The A9 subpopulation of DAn, expressing GIRK2, is required for motor recovery and is preferentially lost in PD ^3,36^. As previously reported for GDNF ^18^, exposure to SEMA3A, WNT5A or NTN1 increased the proportion of GIRK2^+^ A9-like DAn compared to control grafts (Control: 29.50% ± 2.95%; GDNF: 50.40% ± 2.65%; SEMA3A: 58.69% ± 3.90%; WNT5A: 59.23% ± 4.94%; and NTN1: 63.81% ± 1.69%), Figure 2S,T.

**Figure 2.**
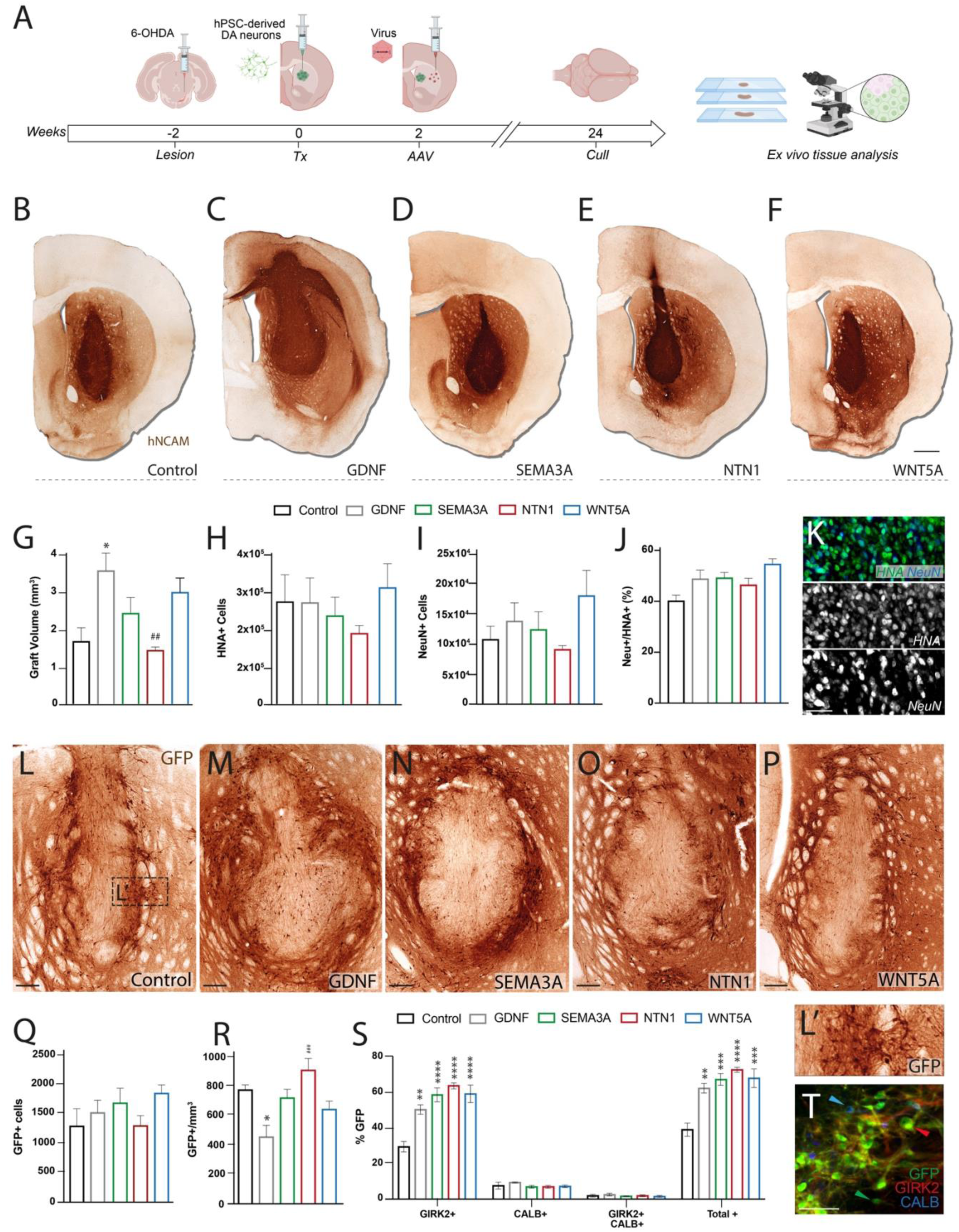
Viral delivery of NTN1, SEMA3A or WNT5A promotes A9 mDA subtype specification. (A) Schematic showing the study design inclusive of 6-OHDA lesioning, neural transplantation, AAVs delivery and histological analyses in mice. (B-F) Representative bright images of VM progenitors grafts revealed by human specific neural cell adhesion molecule (hNCAM) staining in animals that received (B) AAV-empty vector/Control, (C) AAV-GDNF, (D) AAV-SEMA3A, (E) AAV-NTN1 or (F) AAV-WNT5A. (G) Quantification of graft volume, (H) total HNA^+^ cells, (I) total NeuN^+^ cells and (J) proportion of NeuN^+^ cells. (K) Representative images showing the density of cells (HNA^+^) and neurons (NeuN^+^) within a graft. (K) Representative image showing cells within a graft immunolabeled with HNA and NeuN. (L-P) Example bright field images of dopaminergic (DA) neurons, identified by GFP immunostaining, showing similar number of GFP^+^ cells in animals that were exposed to (L) AAV/Control, (M) AAV-GDNF, (N) AAV-SEMA3A, (O) AAV-NTN1 or (P) AAV-WNT5A. (L*’*) Representative image of GFP^+^ cells within a control graft, illustrating cell morphology. (Q) Quantification of total number of GFP^+^ (DA) neurons and (R) density of GFP^+^ neurons. (S) Quantification and (T) representative pictures showing that NTN1, WNT5A and SEMA3A grafts contained a significantly greater proportion of GFP^+^ cells maturing into GIRK2-expressing rather than Calbindin expressing neurons, reflective of A9-like and A10-like DA neurons, respectively – as previously observed in GDNF treated grafts. Data expressed as mean *±*SEM, n=4-5. (G-J, Q-S) One-way ANOVA with Tukey*’*s multiple comparisons test. *p<0.05, **p<0.01, ***p<0.001 and ****p<0.0001 comparing to control; ^##^p<0.01, ^###^p<0.001 comparing to GDNF. Scale bars: (B-F) 1 mm; (K) 50 μm; (L-P) 200 μm; (T) 50 μm. Abbreviations: 6-OHDA, 6-hydroxydopamine; AAV, adeno-associated virus; CTRL, control; GDNF, glial cell-derived neurotrophic factor; GFP, green fluorescent protein; hPSC, human pluripotent stem cells; HNA, human nuclear antigen; NCAM, neural cell adhesion molecule; NeuN, Neuronal Nuclei; NTN, netrin; SEMA, semaphorin.

### SEMA3A, WNT5A, or NTN1 overexpression selectively promote striatal innervation by hiPSC-derived neural grafts

To assess the plasticity of DAn we utilised PITX3-GFP immunostaining to discriminate graft-from host-derived DA fibres. All animals showed robust innervation of the striatum (Figure 3A-B). While overexpression of all the guidance proteins promoted significantly elevated innervation density within the dorsolateral striatum at the level of the graft (0.7mm rostral to Bregma), at more protracted distances (3.42mm caudal to the site of cell implantation), GFP^+^ innervation density was significantly increased in those grafts exposed to NTN1 and WNT5A (Control: 15.60% ± 3.22% GFP-immunoreactive pixels; GDNF: 14.45% ± 3.24%; SEMA3A: 33.40% ± 13.52%; NTN1: 60.01% ± 4.16%; WNT5A: 47.09% ± 4.84%), Figure 3C, E.

**Figure 3.**
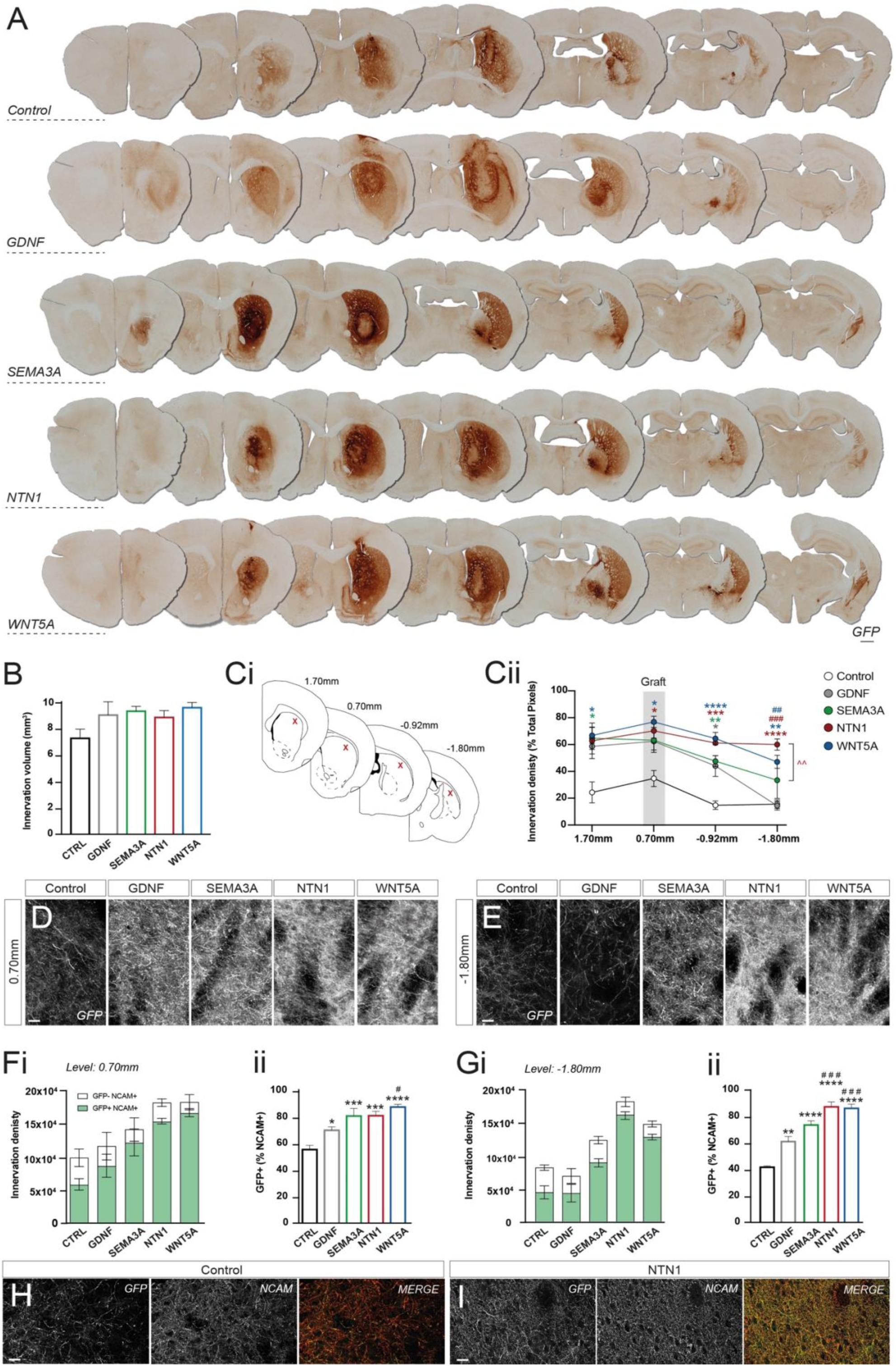
Exposure of hPSC-derived mDA grafts to SEMA3A, NTN1 and WNT5A drives robust innervation of the host striatum. (A) Photomontage of representative VM progenitor grafts highlighting GFP^+^ DA striatal innervation in Control, GDNF, SEMA3A, NTN1 and WNT5A exposed animals. (B) Volume of GFP innervation was similar between all the animals, yet (Ci) sampling of the dorsolateral striatum across the rostrocaudal axis of the brain reveal (Cii) GFP^+^ innervation density was significantly elevated at the site of implantation for all trophin exposed grafts, yet only within the caudal striatum in grafts exposed to NTN1 and WNT5A. (D-E) High magnification of GFP^+^ fibers in the striatum images at the level of the (D) graft core and (E) at distal sites away from cell implantation (−1.8 mm caudal to Bregma, 2.5mm caudal to implantation site). (F-G) Quantification of (Fi,Gi) mDA (GFP^+^/NCAM^+^, green bars) or non-mDA (GFP^-^/NCAM^+^, white bars) graft-derived fibers and (Fii,Gii) the proportion of GFP^+^ fibers at differing rostrocaudal sites in the dorsolateral striatum. GDNF, SEMA3A, NTN1 and WNT5A promoted significantly higher GFP^+^ mDA fibers density and proportions, with NTN1 and WNT5A exposed grafts resulting in superior DA plasticity compared to GDNF, most notably at caudal levels of the striatum. (H-I) Representative examples of immunohistochemistry for GFP (green) and NCAM (red) enabled discrimination between graft-derived mDA and total fiber patterns, respectively. (H) Control grafts had significantly lower GFP^+^ mDA fibers density compared to (I) NTN1. Data expressed as mean ± SEM, n=4-5. (B, C, F, G) One-way ANOVA with Tukey’s multiple comparisons test. *p<0.05, **p<0.01, ***p<0.001 and ****p<0.0001 comparing to control; ^#^p<0.05, ^##^p<0.01, ^###^p<0.001 comparing to GDNF; ^^ p<0.01 comparing SEMA3A. Scale bars: (A) 1 mm; (D,E,H,I) 20 μm. Abbreviations: CTRL, control; GDNF, glial cell-derived neurotrophic factor; GFP, green fluorescent protein; NCAM, neural cell adhesion molecule; NTN, netrin; SEMA, semaphorin.

We next directly assessed DA and non-DA innervation within the host striatum in response to the trophic cues. Human-specific polysialylated neural cell adhesion molecule (hNCAM) and GFP enabled clear discrimination between graft-derived DA (NCAM^+^/GFP^+^) and non-DA (NCAM^+^/GFP^-^) fibres (Figure 3F-H). At the level of the graft core, overexpression of all axon guidance cues significantly increased the relative proportion of DA to non-DA innervation compared to control, (Control: 57.58 % ± 3.16% GFP^+^; GDNF: 72.73% ± 2.57%; SEMA3A: 83.95% ± 5.84%, NTN1: 84.21% ± 3.35%; and WNT5A: 91.02% ± 1.99%), while only WNT5A exposed grafts showed DA innervation levels significantly greater than GDNF (Figure 3Fi-ii). At caudal regions of the striatum (−1.80 mm posterior to Bregma), both NTN1 and WNT5A showed a significantly higher percentage of GFP^+^ DA fibres, compared to control or GDNF (Control: 43.51% ± 0.95%; GDNF: 63.75% ± 3.90%; NTN1: 87.13% ± 3.75%; and WNT5A: 86.89% ± 2.26%) (Figure 3Gii, H, I). Taken together, the selected DA-specific axonal guidance cues positively impacted the plasticity of DAn innervating the striatum, with WNT5A and NTN1 showing influences greater than GDNF.

### Extra striatal graft innervation is increased by GDNF but not by DA-specific axon guidance treatment

Whilst A9 DAn preferentially target the striatum to control motor function, A10-like DAn within hPSC-derived grafts synapse with other extrastriatal targets to modulate a raft of non-motor behaviours ^37^, with evidence that GDNF exacerbates plasticity of these neurons ^18,19^. Here we assessed the impact of the selected trophic cues on extra-striatal targets including the motor cortex, perirhinal cortex, septum, amygdala, olfactory tubercle and substantia nigra pars reticulata (see schematic inserts in Figure 4A).

**Figure 4.**
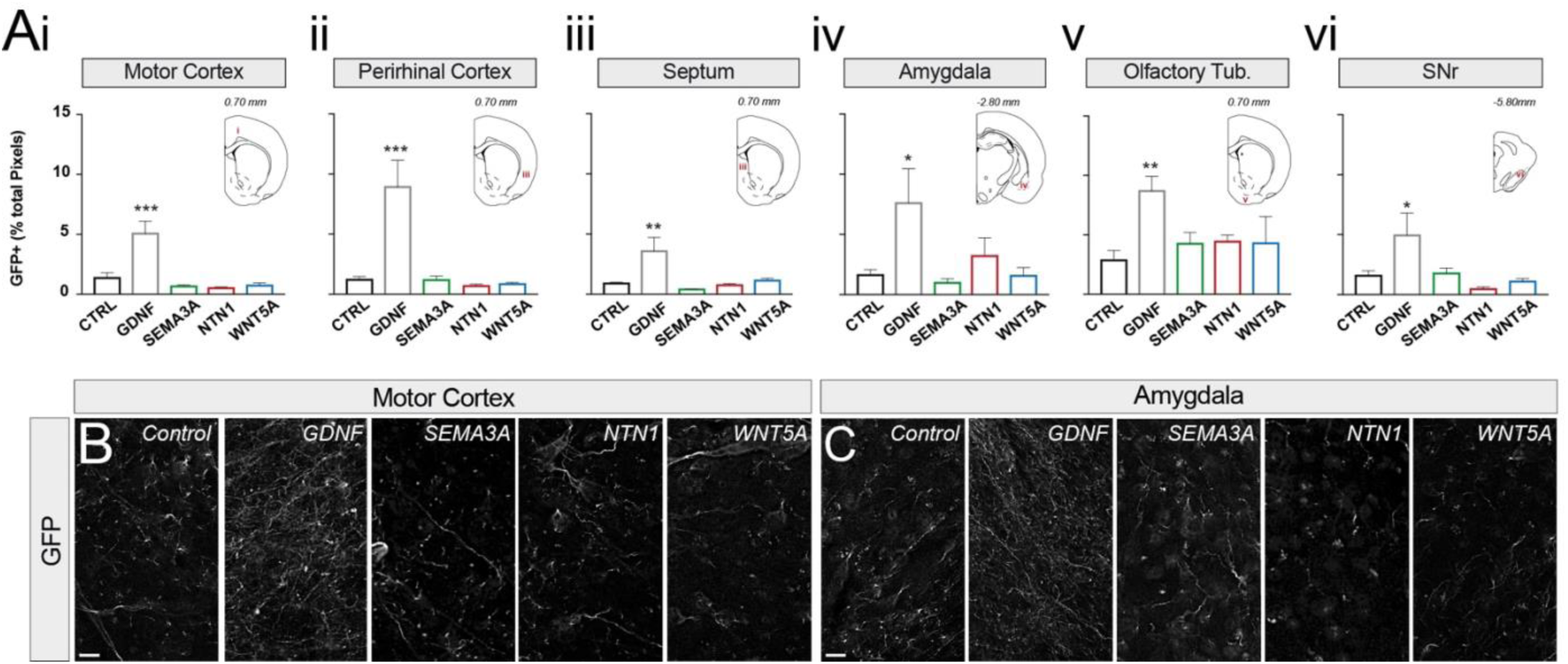
SEMA3A, NTN1 and WNT5A do not promote graft innervation of host nuclei outside the nigro-striatal pathway. (A) Quantification of graft-derived vmDA (GFP^+^) fibers in multiple brain nuclei revealed that only GDNF, significantly increased host innervation outside the striatum. (B) Representative high-power images show that GDNF promotes DA fiber density in the host motor cortex and (C) amygdala. Data expressed as mean ± SEM, n=4-5. (A-C) One-way ANOVA with Tukey’s multiple comparisons test. *p<0.05, **p<0.01, ***p<0.001 and ****p<0.0001 comparing to control; ^#^p<0.05, ^##^p<0.01, ^###^p<0.001, ^####^p<0.0001 comparing to GDNF. Scale bar: 20 μm. Abbreviations: CTRL, control; GDNF, glial cell-derived neurotrophic factor; GFP, green fluorescent protein; NCAM, neural cell adhesion molecule; NTN, netrin; Olfactory Tub, Olfactory Tubercle; SEMA, semaphorin; SNr, substantia nigra pars reticulata.

Graft exposure to GDNF increased GFP^+^ DA innervation across all extra-striatal targets including the motor cortex (GFP fibres = 3.7-fold, compared to control; Figure 4Ai,B), perirhinal cortex (7.9-fold; Figure 4Aii), septum (4.0-fold; Figure 4Aiii), amygdala (4.6-fold; Figure 4Aiv,C), olfactory tubercle (3.0-fold; Figure 4Av) and substantia nigra pars reticulata (3.1-fold, Figure 4Avi), confirming previous observation that GDNF elicited effects broadly on not only A9 DAn but additionally on A10-like DAn ^18,19^. By comparison, graft exposure to the “DA-selective” axonal guidance cues (identified from the snRNA sequencing screen inclusive of NTN1, SEMA3A and WNT5A), revealed GFP fibre density comparable to the control, Figure 4A,B.

### NTN1 overexpression enhances motor recovery in Parkinsonian rats

To evaluate the functional impact of a DA-specific neurotrophic cue in a rat Parkinsonian model, we selected NTN1 for further investigation and compared it to GDNF, a factor consistently shown to improve motor recovery in hPSC-derived transplants ^18–20^. We chose NTN1 due to the discrete graft cores (resulting in reduced host tissue striatal tissue displacement), high DA neuronal numbers, high proportion of A9-like DA neurons (Figure 2), consistent striatal innervation throughout the rostrocaudal axis of the target nuclei (Figure 3), and a reduced proportion of extrastriatal innervation when compared to GDNF (Figure 4).

Two weeks after 6-OHDA lesioning, only rats displaying motor deficits (>300 rotations/hr following amphetamine-induced rotational testing), accounting for 28 of 30 6-OHDA lesioned rats, were stratified into 4 groups: (i) Lesion, Lx; (ii) Lesion + transplant, Tx; (iii) Lesion + Transplant + GDNF-AAV, Tx+GDNF; (iv) Lesion + Transplant + Netrin1-AAV, Tx+NTN1. Rats received intrastriatal grafts of PITX3-GFP vm progenitors, followed by AAV5 delivery (GDNF or NTN1) 2 weeks post-transplantation (Figure 5A). The 6-OHDA-induced motor deficit was persistent and stable over the duration of the study (24 weeks) (Lx, Figure 5B, black symbols), with complete functional recovery in amphetamine-induced rotation observed by 16 weeks in animals receiving grafts in the presence of GDNF or NTN1 (grey and red symbols, respectively) and transplant only (Tx) animal at 20 weeks (Tx, white symbols, Figure 5B). In the adjusted stepping task, no improvement was observed in animals receiving transplants alone (Control: 7.16 *±*2.43% touches with the contralateral paw, relative to the ipsilateral paw; Tx: 5.26% + 1.41). In contrast, a 2.1-fold increase in adjusted steps was observed in NTN1 exposed grafts (14.9% + 2.9, red bar, p=0.47) relative to the Lesion group, reaching significance only in animals receiving grafts exposed to GDNF (25.6 + 5.41, grey bar, Figure 5C). No difference was observed between GDNF and NTN1.

**Figure 5.**
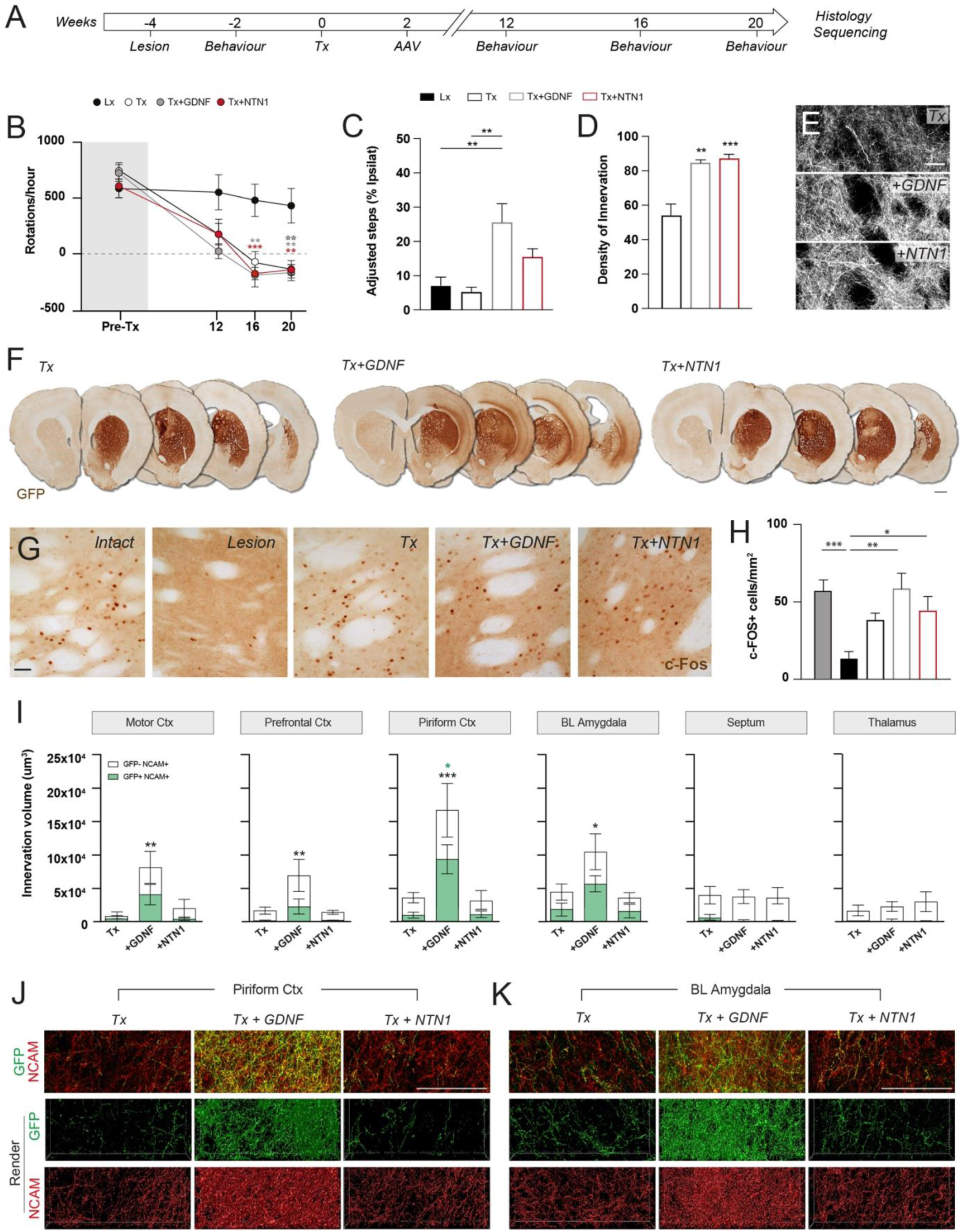
NTN1 improves motor function yet, in contrast to GDNF, shows no evidence of plasticity of non-dopaminergic neurons. (A) Schematic study design inclusive of 6-OHDA lesioning, transplantation ± AVV delivery, and behavioural testing in PD rats with terminal histological and snSEQ assessment of grafts. (B) Unilateral 6-OHDA-lesioned rats showed sustained rotational asymmetry upon amphetamine challenge (black symbols), that could be reversed by 16 weeks in animals receiving vm progenitors grafts exposed to GDNF (grey) or NTN1 (red). Full recovery was observed at 20 weeks in Transplant only (Tx, open circles) animals. (C) NTN1 exposed grafts resulted in a 2.2-fold improvement in the adjusted stepping task, reaching a significant improvement only in GDNF exposed grafts. (D) Reflective of improved motor function, NTN1 and GNDF exposed grafts resulted in a significant increase in the density of GFP^+^ DA innervation in the host dorsolateral striatum. (E) Representative images depicting GFP+ fiber density in the dorsolateral striatum. (F) Representative images showing a PSC-derived vmDA progenitor graft (Tx) and in the presence of GDNF (Tx + GDNF) or NTN1 (Tx + NTN1) in a PD rat model. Note the increased extrastriatal GFP innervation in GDNF exposed grafts. (G) Representative c-Fos^+^ cells in the host striatum, and (H) quantification highlighting that both GDNF and NTN1 significantly increased the number of postsynaptic cells activated by the DA-rich graft. (I) Quantification of graft-derived dopaminergic (GFP^+^, Green bars) and non-dopaminergic (GFP^-^NCAM^+^, white bars) innervation of multiple extrastriatal nuclei, involved in numerous motor and non-motor functions, revealed exposure to GDNF increased off-target innervation. Black and green asterisks represent significant differences in NCAM and GFP, respectively. (J) Representative photomicrographs depicting graft-derived GFP and hNCAM staining in the host piriform cortex and basolateral amygdala, and corresponding renders utilized to quantify innervation density across the graft groups. Data expressed as mean ± SEM, n=5-7/group. One-way ANOVA with Tukey’s multiple comparisons test. *p<0.05, **p<0.01 and ***p<0.001 comparing to Tx. Scale bar: (E, G, J) 100μm; (F) 1mm. Abbreviations: BL, basolateral; Ctx, cortex; GDNF, glial cell-derived neurotrophic factor; GFP, green fluorescent protein; NCAM, neural cell adhesion molecule; NTN, netrin; Olfactory Tub, Olfactory Tubercle; SEMA, semaphorin; SNr, substantia nigra pars reticulata; Tx, Transplant.

At 20 weeks post-transplantation, all animals showed surviving grafts, visible by NCAM and GFP expression within the host striatum (Figure 5F). Where relevant, immunostaining for GDNF or RFP confirmed expression of GDNF or NTN1 protein (data not shown). Confirming observations in mice (Figure 3), exposure of hiPSC-derived DAn grafts to NTN1 or GDNF resulted in a significant increase in the density of GFP^+^ DA fibres within the dorsolateral striatum (Figure 5D,E). To further validate that increased GFP^+^ innervation in the host reflected enhanced integration of the graft into the host circuitry we assessed c-Fos labelling in the striatum of animals injected with D-amphetamine 1 h prior to perfusion. C-Fos^+^ quantification was performed within locations corresponding to dense GFP^+^ fibre density, and targets underpinning motor function – in the dorsolateral striatum (Figure 5G). As anticipated, 6-OHDA lesioning reduced c-Fos expression within the striatum compared to the intact brain (Figure 5H). While notable increases in c-Fos^+^ cells were observed in the dorsolateral striatum of transplanted animals (Tx), only grafts exposed to GDNF or NTN1 resulted in significantly increased c-Fos^+^ density compared to the Lesion (Lx), with restoration comparable to that of the intact brain.

With verified GDNF and NTN1 impacts on graft-induced recovery of motor function validated by *‘*on-target *’* striatal innervation by vmDA neurons and activation of post-synaptic medium spiny striatal neurons, we finally assessed the selectivity of the trophic cues – examining DA (GFP^+^hNCAM^+^) and non-DA (GFP^-^hNCAM^+^) plasticity in off-target (non-striatal) nuclei that encompass a broad range of functions. Regions of assessment included the motor cortex (voluntary movement), prefrontal cortex (responsible for executive functioning), piriform cortex (central in the detection and discrimination of olfactory stimuli), basolateral amygdala (critical for processing emotion and reward-based learning), septum (emotion regulation, reward, and motivation) and thalamus (necessary for relaying sensory and motor information to the cortex, as well as controlling consciousness, sleep and alertness).

Compared to Control (Tx), the density of non-dopaminergic (GFP^-^hNCAM^+^) fibres emanating from the graft was significantly higher for those grafts exposed to GDNF within areas including the motor (+8.62-fold), prefrontal (+4.20-fold) and piriform cortex (+4.56-fold) as well as basolateral amygdala (+2.37-fold), Figure 5I-K. The density of GFP^+^ DA fibres was also significantly increased within the piriform cortex from GDNF-exposed grafts, similar to observations in the mice (Figure 4), and trending increases also in the motor cortex and amygdala. Highlighting selectivity, no increase in innervation density of these targets was observed for grafts exposed to NTN1, Figure 5I.

### NTN1 is a DA-selective neurotrophic and axon guidance cue

To verify the observed specificity of NTN1 for DA neurons we performed single nuclei sequencing on these same hPSC-derived grafts exposed to NTN1 or GDNF, using adjacent serial sections. Three grafts were pooled for each of the graft conditions (i.e. Tx, Tx + GDNF and Tx + NTN1). Human nuclei were scored using gene sets representing human or rat signature genes (Figure 6A, Supplementary Figure 4B). The subsets of nuclei denoted as *“*human” across the samples were subsequently merged, integrated, normalized and clustered. Cluster analysis revealed 16 populations that were confirmed to be comparatively represented across the samples (Supplementary Figure 4C). Based on key markers, clusters were defined as Neurons (SYP, SYN1, GAP43, MAPT), astrocytes (GFAP, AQP4), oligodendrocytes (OLIG1), neural stem cells (SOX2) or mix (Figure 6B, Supplementary Figure 4D). Neurons were subsequently clustered as DA or non-DA based upon key marker expression (TH, EN1, DRD2, KCNJ6) (Figure 6C, Supplementary Figure 4F).

**Figure 6.**
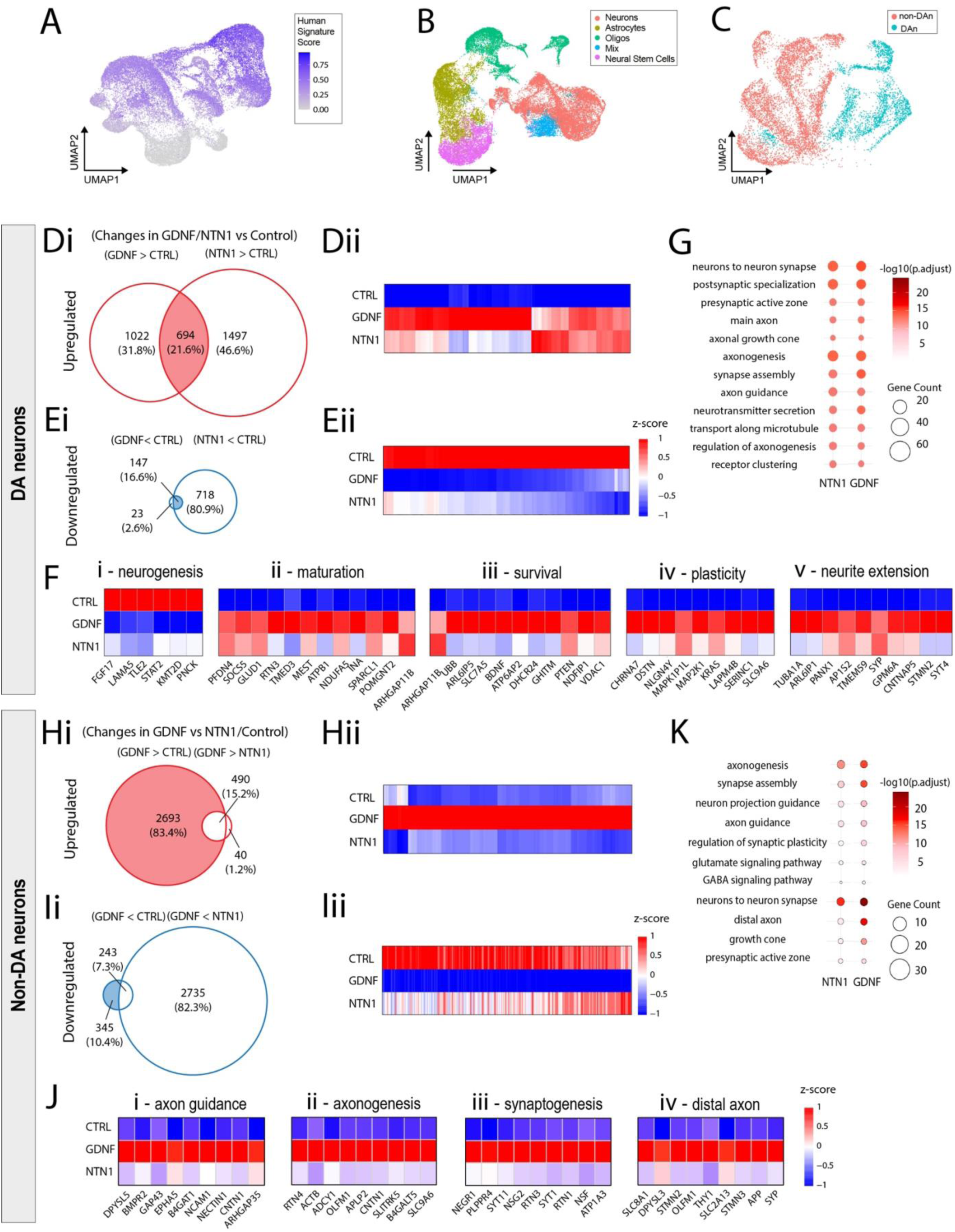
Transcriptional profiling of NTN1 exposed vm grafts confirm the specificity for DA neuron plasticity and not other neuronal populations. (A) UMAP showing how the human nuclei were identified by analysis of the relative proportion of mRNA transcripts mapping to the human transcriptome. (B) Human nuclei based on initial Louvain clustering were annotated as neurons, astrocytes, oligodendrocytes, neural stem cells and a *‘*Mix*’* of other indiscernible cell types. (C) The neuronal population was identifiable as DA (cyan) or other neurons (pink). (D) Venn diagram showing differentially expressed genes within the grafted dopamine neurons in response to GDNF and NTN1, highlighting overlapping upregulated and downregulated genes. (E) Down regulated genes in DA neurons in response to GDNF and NTN1 included those associated with neurogenic processes (Ei), whilst upregulated genes were linked to neuronal maturation, survival, plasticity and axon morphogenesis (Eii-v). (F) Gene ontology highlighting similarly upregulated gene clusters in response grafted DA neurons in response to GDNF and NTN1. (G) Venn diagram showing differentially expressed genes between GDNF and NTN1 exposed grafts and heat maps highlighting differentially down and upregulated genes in response to GDNF that remained no significantly changed in control or NTN1. (H) Many of the differentially upregulated genes in non-DA neurons were associated with axonal plasticity and morphogenic processes including axon guidance, axongenesis, synaptogenesis and distal axon. (I) *‘*GO *’*terms identifying clusters of genes associated with GABA and glutamatergic transmission and/or plasticity in non-DA neurons in response to GDNF but not NTN, highlighting the lack of selectivity of GDNF. Abbreviations: CTRL, control; DA, dopamine; GDNF, glial cell-derived neurotrophic factor; NTN, netrin.

We initially focused on the DA neuron population and performed differential gene expression analyses, comparing the GNDF and NTN1 exposed grafts to Control. This revealed a total of 694 DEGs upregulated and 147 DEGs downregulated in response to both GDNF and NTN1 compared to control, (Figure 6D,E – refers to shaded areas in venn diagrams). Numerous downregulated genes (Log2-fold change, Figure 6D), corresponded to those involved in neurogenic and cell proliferation processes such as the mitogen fibroblast growth factor 17 (FGF17), the basement membrane protein laminin A5 (LAMA5), essential in neural tube formation and Transducin-Like Enhance (TLE2), a corepressor that inhibits neuronal differentiation (Figure 6Gi). Complementary, was the upregulation of pro-neural and maturation genes including neuronal intermediate filament/microtubule-associated proteins PFDN4 and INA, the JAK-STAT signaling regulator Suppressor of Cytokine Signaling 5 (SOCS5) and the vesicular trafficking receptor Transmembrane P24 Trafficking Protein 3 (TMED3) (Figure 6Fii). Supporting known roles for GDNF and NTN1 in neuronal survival was the expression of genes including SLC7A5, PTEN and NDFIP1 (all involved directly or indirectly in the PI3K-Akt pro-survival pathway), Figure 6Fiii. Finally, reflective of the increase dopaminergic innervation observed from GDNF or NTN1-exposed grafts were genes associated with plasticity, axon growth/guidance and synaptogenesis including MAP2K1, MAPK1P1L, SYT4, SYP and SV2B (Figure 6Fiv-v). Gene ontology analysis of biological processes and cellular compartments further highlighted the remarkably similar profiles between the grafted dopamine neurons exposed to NTN1 or GDNF, compared to controls – showing significant upregulation of gene clusters associated with *‘*neuron to neuron synapse*’*, *‘*presynaptic active zone*’*, *‘*axonogenesis*’*, *‘*neurotransmitter secretion and *‘*axon guidance *’*(Figure 6G).

Delving further into the histological observation of GDNF promoting plasticity in non-DAn (Figure 5I,J), we subsequently performed differential gene analysis on non-DAn exposed to GDNF and NTN1. Comparative analysis revealed significant differences. Of the 3183 DEGs upregulated in response to GDNF compared to control, 2693 were uniquely upregulated after GDNF but not NTN1 exposure, Figure 6H. Conversely, of the 588 downregulated genes in response to GDNF, 345 were unique with no significant change in response to NTN1 (Figure 6I, see blue shaded area in venn diagram). Similar to DA neurons, these downregulated genes were implicated in neurogenic and cell proliferation pathways. Most noteworthy was the upregulation of plasticity-associated genes (including axon guidance, axogenesis, synaptogenesis and distal axon) in response uniquely to GDNF, yet not NTN1, inclusive of GAP43, NCAM, CNTN1, SLITRK, SYT11, SYT1, SYP (Figure 6J).

Verifying the lack of specificity of GDNF for non-DA neurons, was the significant upregulation of Biological Processes (BP) gene ontology terms associated with axogenesis, synapse assembly and axon guidance, inclusive of glutamate and GABA associated signaling. Cellular Component GO terms significantly upregulated in non-DA neurons in response to GDNF, but not NTN1, included *‘*neuron to neuron synapse*’*, *‘*distal axon*’*, growth cone *’*and *‘*presynaptic active zone *’*(Figure 6K).

Taken together this data highlights the specificity of NTN1 for DA neurons, in contrast to GDNF, which acts as a potent neurotrophic cue on both DA and non-DA neuron populations.

## DISCUSSION

With several Phase I clinical trials examining the safety of human pluripotent stem cell-derived neural grafts for the treatment of PD coming to fruition ^38–40^ or ongoing (see review ^41^), the task ahead will be deciphering the full functional benefits of these transplants. Whilst current and planned trials focus on the donor material ^41,42^, preclinical evidence suggests that modulation of the host environment will also likely have a significant impact in improving the ability of these transplants to restore function, see review ^30^. Most relevant have been findings showing that prolonged GDNF exposure to hPSC-derived grafts (delivered virally or via biomaterials) ^18–20^ improve graft survival, integration and recovery of motor deficits in rodent models of PD. However, GDNF lacks specificity for DA neurons and the implications of non-DA plasticity on graft and host function remains unknown. Here, we performed single-nuclei RNA-sequencing of maturing hPSC-derived VM neural grafts to identify DA-selective axon guidance cues that may be exploited to more selectively drive DA plasticity and functional integration whilst reducing off-target innervation and their potential associated effects. Single nuclei analysis of DA neurons within the grafts showed upregulation of numerous molecules associated with the WNT, Netrin and Semaphorin signalling pathways that can be driven by WNT5A, NTN1 and SEMA3A respectively, and have been shown to support DA fibre extension during the development of neural circuits, including the nigrostriatal pathway *in vitro* and *in vivo* (for reviews see ^26,43^). In rodent models, the selectivity of these trophic cues has previously been reported. *In vitro*, NTN1 has been shown to enhance DA neurite outgrowth ^30,44^, and its overexpression prevents lesion-induced degeneration of nigral DA neurons, as well as promoting DA innervation and functional recovery ^35^. Similarly, WNT5A promotes plasticity of rodent midbrain DA neurons *in vitro* and in fetal tissue grafts ^27,29^, whilst SEMA3A stimulates DA nigro-striatal axonal pathfinding in development and induces DA plasticity of VM DA neurons derived from fetal tissue ^45^ and pluripotent stem cells ^28^.

Overexpression of each of the selected guidance cues (SEMA3A, WNT5A and NTN1) significantly improved neural graft outcomes in a rodent Parkinsonian model. All trophic cues tested significantly increased the proportion of A9-like neurons, compared to the control, with trends of greatest impact in grafts exposed to NTN1. This A9 subpopulation of dopamine neurons is responsible for control of motor function ^36,46^ including the reversal of motor dysfunction in preclinical neural grafting studies in Parkinsonian rodents ^3,47^. Recent findings have shown that the acquisition of A9 and A10-like phenotypic specification of vm DAn with hPSC-derived neural grafts is underpinned by striatal versus cortical synaptic target acquisition ^2^. Reflective of this, grafts exposed to SEMA3A, WNT5A and NTN1 showed significantly increased DA (GFP^+^) innervation of the striatum compared to the control, with both NTN1 and WNT5A elevating DA innervation across the entire rostro-caudal axis of the striatum, and to levels significantly greater than GDNF. Of note, NTN1 exposed grafts were smallest in core size compared to other treatments, yet showed maximal striatal innervation, indicative of high plasticity of these DA neurons compared to all other graft groups.

Furthermore, the degree of DA plasticity evoked by SEMA3A, NTN1 and WNT5A exposure was notably selective for A9 DAn, with only marginal DA innervation in extrastriatal targets. In contrast, exposure of grafted DAn to GDNF resulted in high level of ‘off-target’ DA plasticity in the motor and perirhinal cortex, septum, amygdala, olfactory tubercle and substantia nigra pars reticulata - a collection of target nuclei innervated by A10 VTA DA neurons that underpin behaviours inclusive of sensory processing, memory consolidation and aggression. Future studies, such as fluorogold back-tracing, will be required to further untangle the ligand-receptor interactions that drive these degrees of selectivity between dopaminergic neuron subtypes and across striatal compartments and broader regions of the brain. Unravelling these complexities will be influenced by our knowledge of development, where key examples suggest that downregulation of known axon guidance genes may just as importantly modulate responsiveness of grafted neurons - noting for example that a reduction/depletion of UNC5 can change the responsiveness to NTN1 from repulsion to promoting growth of vmDA axons towards their rostral targets ^48–50^. In addition is the example of defined Plexin-Semaphorin interactions that play key roles in separating the VTA and SNc axons, through growth inhibitory mechanisms, during establishment of their forebrain trajectories ^51^.

Our ability to more precisely interrogate dopamine and non-dopamine neurons within the hPSC-derived vm neural grafts, using single nuclear sequencing on fixed tissue (from animals examined in parallel behavioural and histological assessments) enabled clear scrutiny of the selectivity of NTN1. While DEG profiles were largely comparable for the DA neurons within grafts exposed to NTN1 or GDNF, the non-DA neurons exposed to GDNF showed high transcriptional changes in genes and pathways associated with axonal plasticity, guidance and synaptogenesis – reflective of the off-target effects of GDNF compared to NTN1. Whilst the present findings highlight the specificity of an alternative guidance cue to drive (A9) DA plasticity and reduce ‘off-target’ DA and non-DA plasticity, these observations need to remain in check with the goals of reversing motor deficits. Here we report comparable motor recovery between GDNF and NTN1 exposed grafts in a pharmacologically induced task (amphetamine-induced rotational testing) yet show only GDNF-exposed grafts enhanced recovery in the non-pharmacological induced adjusted stepping task. Considering the smaller graft size in response to NTN1 compared to GDNF, it remains to be determined whether increase donor cells implanted could comparatively restore these deficits, and/or whether seemingly ‘off-target’ innervation from presumable A10 DA, and/or non-DA neurons also contributes to functional outcomes. Prior rodent studies have clearly highlighted the necessary presence of A9 DAn within grafts to restore motor function ^3^, yet the anatomical organisation, protein expression, projections and thereby classification of human midbrain DAn is notably different ^26^. This suggests that future studies, aimed at silencing defined populations within grafts exposed to GDNF (i.e. the non-DAn and/or A10 DAn), will be necessary to shed more light on the involvement of other human graft-derived populations in restoring motor function.

Additional to this will be the requirement for further studies to understand regulation of gene and consequently protein expression, with evidence that A9/SNc DAn respond to lower concentrations of Netrin than A10/VTA neurons in the developing rodent brain ^52^. Further, will be the need to understand the temporal requirements of NTN1, with evidence of NTN1 not only influencing plasticity and pathfinding having a role of neuroprotection in PD models ^35^. Hence earlier or biphasic delivery, as proposed for GDNF ^18^, may be optimal.

Here we report on guidance cues NTN1, SEMA3A and WNT5A for their benefit in enhancing the structural and functional integration of hPSC-derived DAn in grafts. However, other more specific cues may exist - noting the inherent limitation in single nuclei sequencing approach adopted here that captures only nucelar mRNA hence reflects current transcriptional state and not mRNA outside the nucleus (i.e. in the dendrites, axon growth cone, pre and post-synaptic compartments) ^53^ and thereby may have missed additional genes engaged in axonal pathfinding in our initial screen. Evolving spatial transcriptomic technologies are anticipated to enable detection of whole transcriptome in situ with subcellular resolution, without the need of tissue dissociation. Whilst this has recently been performed on mature hPSC-derived neural grafts, the level of resolution remained below beyond that of the sn/scRNA-seq^54^.

Also worthy of consideration is the observation that GFRα1 was identified in the original single nuclei screen of DA neurons compared to non-DA. The data presented here suggests sufficient expression of the GDNF receptor in the non-DA population to induce trophic responses. By comparison, the differential expression of other axon guidance cues/pathways (i.e. NTN1) was likely at levels that impacted the DA but not non-DA neurons thereby culminating in the select DA over non-DA responsiveness. The findings here highlight the selectivity of NTN1 for A9-like DA neurons.

While we continue to interrogate and understand the nuances of these axon guidance cues and their potential for positively influencing the structural and functional integration of neural grafts for the treatment of PD, here we provide a new piece to the puzzle. We demonstrate increased selective striatal targeting of DA axons, reduced off-target plasticity, whilst retaining the capacity to reverse motor symptoms in Parkinsonian rats, comparable to that recently achieved for GDNF exposed grafts. Future studies may further unravel additional and/or alternative guidance cues that positively influence not only DA plasticity in the quest to improve motor function, but additionally regulate other graft-derived circuits that may impact restoration of cognition or other affected behaviours in PD. In all, these findings hold important implications as the field continues to embark on clinical trials in PD patients.

## Resource Availability

### Lead Contact

Requests for further information and resources should be directed to and will be fulfilled by the lead contact, Clare Parish.

### Materials Availability

Viral vectors generated for this study will be made available upon request to lead contact, Clare Parish.

### Data and Code Availability

The authors declare that the data supporting the findings of this study are presented within the paper and its supplementary information files. Source data will be shared upon request. The code generated in this study is available on github at https://github.com/parishgroup/Axonal_Guidance and https://github.com/zchatt/ASAPSpatialTranscriptomics/blob/main/xenograft/snrna_seq/cellranger.sh

### Author Contributions

CP, CLP conceived the experiments and wrote the manuscript; CP, KHL, CWG, SF, JC, CPJH, BX, EH, ZC, GMH, NM, CLP performed experiments; CLP provided reagents and expertise; all authors reviewed/edited manuscript; CLP provided funding.

## Acknowledgements

The authors thank Dr Yair David Joseph Prawer (The University of Melbourne, Melbourne, Australia) for his support in bioinformatic analyses, Mr David Yoannidis and the Molecular Genomics Core at the Peter MacCallum Cancer Center for their technical support on the library preparation and sequencing. We thank Carlos Gantner for his critical reading of the manuscript. CLP was supported by a National Health and Medical Research Council Australia (NHMRC) Senior Research Fellowship and subsequent NMHRC Leader Fellowship (level 2). NM was supported by a EH Flack fellowship provided by the Marion and E.H. Flack Trust. This work was supported by funding from the NHMRC (GNT2026395 and GNT2038892). The Florey Institute of Neuroscience and Mental Health acknowledges the strong support from the Victorian Government and the funding from the Operational Infrastructure Support Grant.

## STAR METHODS

### Cell culture and differentiation

The male human induced pluripotent stem cell (hiPSC) reporter line, RM3.5 *PITX3*-*eGFP* ^2^, was cultured and differentiated into vm progenitors and neurons as previously described ^55^. In preparation for transplantation, day 19 (19DIV) vm progenitor cultures were dissociated using Accutase (Innovative Cell Technologies) to a single-cell suspension (100,000 cells/μl) in a maturation media comprised of NBB27 (1:1 DMEM/F12 and Neurobasal media, 1x B27 with Vitamin A, 1x N2, 1x Glutamax, 0.5x Penicillin streptomycin, 1x non-essential amino acids and 1x insulin-transferrin-selenium-sodium pyruvate-A) containing brain-derived neurotrophic factor (BDNF, 20ng/ml, R&D Systems), glial cell-line derived neurotrophic factor (GDNF, 20ng/ml, R&D Systems), recombinant human transforming growth factor type β3 (TGFβ3, 1ng/ml, Peprotech), ascorbic acid (200nM, Sigma-Aldrich), dibutyryl cAMP (0.05mM, Tocris), DAPT (10μM, Sigma-Aldrich) and 10μM ROCK-inhibitor Y27632 (Tocris).

### Surgical procedures

All animal procedures were conducted in agreement with the Australian National Health and Medical Research Council’s published Code of Practice for the Use of Animals in Research, and experiments approved by The Florey Institute of Neuroscience and Mental Health Animal Ethics committee. Animals were group housed on a 12:12-hour light/dark cycle with ad libitum access to food and water. Surgery was performed on 28 Swiss mice (Arc:Arc(S)) (to confirm efficiency of the viral vectors), 28 nude mice (BALB/c-Foxn1^nu^/Arc) and 36 athymic nude rats (CBH^rnu^). To model the dopamine depletion associated with Parkinson’s disease, nude rats and mice received a unilateral injection of 6-hydroxydopamine (6-OHDA) into the mouse substantia nigra (2.4ug) or rat medial forebrain bundle (14ug) as previously described ^56^. Two weeks after 6-OHDA lesioning, animals were transplanted with hiPSC-derived *PITX3-eGFP* VM progenitors (100,000 cells in 1μl) ectopically into the striatum and, after a further 2 weeks, animals received unilateral injections into the dorsolateral striatum of adeno-associated viruses (AAV), as previously described ^56^. The AAV vector serotype 5 (0.25 μl) were designed to drive expression of a gene and/or red fluorescent protein (mCherry) *in vivo*. The genes encoded and vectors were as follows: GDNF (AAV5-GDNF, 4.2e^12^ gc/m), SEMA3A (AAV5-SEMA3A, 2.8×10^12^ gc/m), NTN1-mCherry (AAV5-NTN1-mCherry, 3.09×10^12^ gc/m), WNT5A-mCherry (AAV5-WNT5A-mCherry, 2.12×10^12^) or mCherry only (AAV5-mCherry, 4.1e^12^ gc/m), all expressed under the chicken ß-actin (CBA) promoter. See Supplementary Table 1 for virus constructs.

### Behaviour

Two weeks after lesioning rats, motor function was assessed using the amphetamine-induced rotation test. In brief, net rotations over 60 minutes were analysed 10 minutes after intraperitoneal injection of D-amphetamine sulfate (5mg/kg; Tocris Bioscience). Upon completion of initial testing 2 weeks post-lesioning, animals displaying a functional deficit (>300 rotations in 60 min) were ranked in order of the percentage rotational asymmetry and performance matched across the four treatment groups: (i) Lesion only (Lx), (ii) Transplant only (Tx), (iii) Transplant + AVV5-GDNF (Tx+GDNF), (iv) Transplant + AAV5-NTN1 (Tx+NTN1). To assess functional integration of the transplanted cells, rats were re-tested at 12, 16 and 20 weeks after grafting as well as the adjusted stepping test performed at 20 weeks in all animals, as previously described ^56^. All rats received an injection of amphetamine 1hr prior to transcardial perfusion to induce synaptic dopamine release and subsequent activation of the intermediate early response gene, c-Fos, in postsynaptic medium spiny neurons of the striatum, as previously described ^18^.

### Real-Time qPCR

Using quantitative PCR, expression of human SEMA3A, NTN1 and WNT5A were measured from striatal tissue of Swiss mice 1 week after intrastriatal injection of the AAVs. Mice that did not receive any AAV were included as controls. For tissue preparation, mice were culled with an overdose of sodium pentobarbitone (100mg/kg), the striatum (containing the virus) dissected on a chilled plate, snap frozen and kept at −80°C. On the day of analyses, brain samples were homogenized and total RNA was extracted using ISOLATE II RNA Minikit (Bioline, UK), according to manufacturer’s specifications. The RNA was converted to cDNA using the SuperScript VILO cDNA synthesis kit (Invitrogen, USA). Quantitative PCR was performed using the PowerUp SYBR Green Master Mix kit (Applied Biosystems, USA) in an RG-6000 Rotogene system (Corbett Research, Australia). See Supplementary Table 2 for full list of primer sequences utilized in this study.

### Single-nuclei RNA isolation, sequencing and bioinformatic analyses

Brains were rapidly extracted from nude rats after receiving an overdose of sodium pentobarbitone (100mg/kg) and transcardial perfusion with ice-cold PBS. The striatum containing the human iPSC-derived neural graft was dissected on ice, snap-frozen and kept at −80°C. Homogenization and nuclei isolation followed the protocol https://www.protocols.io/view/nuclei-isolation-for-10x-chromium-single-nuclei-rn-cymyxu7w. Briefly, brain samples were homogenized using a glass dounce homogenizer (24 strokes) in 1.5mL of ice-cold lysis buffer (Nuclei EZ Prep, Sigma, with 0.2U/μl recombinant RNase inhibitor, Takara) and incubated on ice for 5 min with 1.5ml of lysis buffer. Centrifuged nuclei (500g, 5 min, 4°C) were washed in 1ml of ice-cold lysis buffer, incubated for 5 min, centrifuged (500g, 5 min, 4°C) and resuspended in 1ml of wash buffer (PBS, Gibco, with 1% Ultrapure Bovine serum albumin, Ambion, and 0.2U/μl RNase inhibitor). Isolated nuclei were filtered through a 30μm cell strainer (MACS SmartStrainers), washed with 1ml of wash buffer, centrifuged (500g, 5 min, 4°C), resuspended in 300μl of wash buffer on ice, counted using Trypan blue (ThermoFisher) and stained with primary antibodies, including human nuclear antigen conjugated to Phycoerythrin (HNA-PE, 1:100, Abcam) and rabbit anti neuronal nuclei (rb-NeuN, 1:100, Abcam). After incubation (30 min, 4°C), rabbit secondary antibody Alexa Fluor 647 (AF647) conjugated was added (1:200, Jackson ImmunoResearch), nuclei suspension incubated (20 min, 4°C, dark) and washed two times with 1ml wash buffer (centrifugation 500g, 5 min, 4°C). Stained nuclei were resuspended in DAPI (1μg/ml, ThermoFisher Scientific) and DAPI^+^ HNA-PE^+^ NeuN-AF647^+^ singlets isolated using FACS ARIA III (BD Biosciences, 70μm nozzle, 21– 22 p.s.i.) - see Supplementary Figure 1C for gating strategy. As negative controls, human hiPSC, mouse only or unstained brain tissue were used (Supplementary Figure 1B). Sorted nuclei were counted twice using automated NuceloCounter ® NC-200, prior to loading into the 10x Chromium (10x Genomics). Library preparation for all samples followed the Chromium Single Cell 3′ Library & Gel Bead Kit v3.1 (CG000315_ChromiumNextGEMSingleCell3-_GeneExpression_v3.1_DualIndex RevB.pdf.) with 13 complementary DNA pre-amplification cycles and sequencing on NovaSeq S1 (Illumina) lane using a custom Chromium run configuration of 28:10:10:90. A Human (GRCh38) and Rat (mRatBN7) reference (GRCh38_and_mRatBN7-2023-A_build) genome was constructed using “cellranger mkref” as outlined in https://github.com/zchatt/ASAP-SpatialTranscriptomics/blob/main/xenograft/snrna_seq/GRCh38_and_mRatBN7-2023-A_build.sh.

Raw FASTQ files were aligned to GRCh38_and_mRatBN7-2023-A_build using “cellranger count” as detailed in https://github.com/zchatt/ASAPSpatialTranscriptomics/blob/main/xenograft/snrna_seq/cellranger.sh.

Single-nuclei data were processed using Seurat ^57^ environment in R programming language. Nuclei counts were normalised using SCTransform. High quality single-nuclei were retained that had < 0.1% mitochondrial RNA and > 2000 features (Supplementary Figure 2A, B). Nuclei in which >90% of mRNA were aligned to human transcripts were denoted as human origin (Supplementary Figure 2C). Human nuclei were then re-clustered (Leiden algorithm). To evaluate the cell-types of the human nuclei we used a reference-based single-cell RNA-seq annotation tool (scType) ^58^. Differential expression analysis was performed using LIMMA Voom using the raw counts within xenograft and SNpc scRNAseq datasets ^59^. Gene ontology enrichment was performed using enrichR ^60^ and GO_Molecular_Function_2015” and “GO_Cellular_Component_2015” databases.

### Single-nuclei RNA isolation, sequencing and bioinformatic analyses <u>fixed samples</u> (PathoSeq)

The protocol for single nuclei isolation of fixed tissue was adapted from Wang et al., 2024 ^61^. In brief, the human vmDA graft was visibly identifiable and dissected using a punch biopsy needle (2mm, KAI-700, 21056-KA) from serial brain slices. Grafts from 3 animals per condition were pooled together in a 1.5ml Eppendorf tube, with 100μl of Digestion buffer (PBS^-/-^ with 1mg/ml Liberase TH (Roche Diagnostics), 0.2U/μl RNase Inhibitor, 0.5mM CaCl_2_) and mechanically homogenized using a Biomasher II (Biostrategies), with 30 strokes. Tissue was then digested in a total of 1ml of digestion buffer (37°C, 45 min, 800rpm) on a Thermomixer, 400μl of EZ buffer added to the sample (EZ Lysis Buffer with 1% BSA, 0.2U/μl of RNase inhibitor), sample mixed by inversion 5x and centrifuged for 5 min at 900g at 4°C. Supernatant was removed and pellet resuspended in 600μl of EZ buffer, mixed 5x by pipetting, incubated 5 min on ice, passed through a 25G needle (5-10x) and washed with 600μl of wash buffer (PBS^-/-^ with 1%BSA and 2.5mM MgCl_2_, 0.2U/μl RNase inhibitor). The tissue was then pelleted by centrifugation (4°C, 900g, 5 min), resuspended in 1000μl of wash buffer gently using a p1000, pipetting 30x, filtered (70μm and 40μm PluriStrainer Mini filter). Single nuclei libraries were constructed using the GEM-X Flex Gene Expression kit for multiplex samples (10x Genomics), following the 10x user guide (CG000787_GEM-X_Flex_MultiplexedSamples_UserGuide_Rev_B). Nuclei were quantified using the LUNA-FL dual fluorescent cell counter and equal numbers of nuclei were combined for each sample. Libraries were sequenced on NextSeq2000 (Illumina) P4 flow cells, 100 cycles. Reads were demultiplexed and mapped to the reference genome (GRCh38-2024-A, build 2024-A, 10x Genomics) using Cell Ranger v9.0.1 (10x Genomics). As above, single-nuclei data were processed using Seurat ^57^ environment in R programming language. For each sample, high quality single-nuclei were retained that had < 10% mitochondrial counts, between 500 and 10000 features and more than 1000 RNA counts (Supplementary Figure 4A). Samples were then normalized, scaled and PCA, UMAP and clustering performed using Seurat inbuilt functions (NormalizeData, ScaleData, RunPCA, RunUMAP, FindNeighbours, FindClusters). Graft and host signature (Supplementary Figure 4B) was derived from the top and bottom of the first principal component of a dataset with known human and rat brain snRNA-seq (see list of genes on github https://github.com/parishgroup/Axonal_Guidance) and the subset of nuclei denoted as *“*human”. Human nuclei from each sample were then merged, integrated, normalized and clustered (Supplementary Figure 4C). Based on key markers, clusters were defined as neurons (SYP, SYN1, GAP43, MAPT), astrocytes (GFAP, AQP4), oligodendrocytes (OLIG1), neural stem cells (SOX2) or mix (Supplementary Figure 4D). Nuclei identified as neurons were retained in a new Seurat object (Supplementary Figure 4E), normalized, scaled and clustered and nuclei were classified as dopaminergic (TH, EN1, DRD2, KCNJ6) or not (Supplementary Figure 4F). Differential expression analysis was performed using Seurat function FindMarkers and gene ontology enrichment using enrichR ^60^ and “GO_Molecular_Function” and “GO_Cellular_Component” databases.

### Immunohistochemistry

Confirmation of VM specification of the PITX3-eGFP hiPSCs was performed at day 13 and 25 of differentiation on paraformaldehyde (PFA, 4%, 10mins) fixed cultures. To assess in vivo graft survival, composition and integration, mice and rats were given an overdose of sodium pentobarbitone (100mg/kg) at 4 or 24 weeks after grafting and were subsequently transcardially perfused with ice-cold PBS followed by 4% PFA. The brains were cryoprotected in 20% sucrose solution and cryosectioned on a freezing microtome (40μM; 12 series). Immunocytochemistry (ICC) was performed on fixed cultures and immunohistochemistry (IHC) on free floating brain sections as previously described ^62^. Hematoxylin and eosin (H&E) staining on 5μm sections, to confirm the presence of a neural graft prior to snRNA sequencing, was performed on perfusion fixed (10% neutral buffered formalin) brains.

### Microscopy and quantification

Brightfield, darkfield and fluorescence images were captured on an Invitrogen EVOS M5000 Imaging System (for cell cultures, 10x/0.30NA and 20x/0.45NA air objective), a Leica DM6000 upright epifluorescent microscope (for volumetric and number of GFP^+^, GIRK2^+^, Calbindin^+^, HNA^+^, NeuN^+^ cells count, 10x/0.30NA and 20x/0.80NA air objectives), and a Leica Thunder inverted microscope (for GFP^+^ and NCAM^+^ fibre analyses, 20x/0.80NA air objective). Where necessary LASX (Leica) software was used to generate multi-tiled images or computational clearing. All optical density measurements were captured in single microscopy sessions to ensure standardisation of optical settings, including light intensity, across all examples used for comparative analysis. Graft volumes (Figures 2G, 3G) and mDA innervation volume were delineated by GFP expression and volume extrapolated using Cavalieri’s principle. For cell counting (Figure 2), all GFP^+^ cells were counted across the rostrocaudal axis of the entire graft in a 1/12 series from brightfield images and corrected for series number; the fraction of GIRK2^+^ and CALB^+^ expressing cells was quantified by examining all GFP^+^ cells across fluorescent immuno-labelled sections containing the graft core; HNA^+^ and NeuN^+^ cells were counted in 3 fields of view (20x) spaced across the axis of the graft and total numbers estimated using density and graft volume. GFP^+^ and NCAM^+^ innervation density (Figures 3, 4) were assessed in a single field of view (FOV, 20x) of 1/12 series of fluorescent immunohistochemistry as previously described ^19^. Briefly, 10 z-stack-sections (1μm per section) were obtained and compressed, NCAM^+^ and GFP^+^ fluorescence channels were separated and NCAM^+^ or GFP^+^ fibres were isolated on colour inverted images using the ‘colour range’ tool on Photoshop (Adobe). GFP^+^ fibres percentage was calculated by dividing GFP^+^ immunoreactive pixels by NCAM^+^ immunoreactive pixels. Innervation density was expressed as number of immunoreactive pixels or percentage of GFP^+^ pixels. All areas were captured with conserved settings. Sampling for fibre density was performed (from 1.7mm anterior to −1.8mm posterior to Bregma) in the dorsolateral striatum, motor cortex, perirhinal cortex, septum, amygdala, olfactory tubercles and olfactory tubercles of grafted mice.

### Statistical Analyses

All data are presented as mean ± SEM. Statistical tests employed (inclusive of one- and two-way ANOVA and student t-tests) are stated in figure legends. Alpha levels of p<0.05 were considered significant with all statistical analysis performed using GraphPad Prism. *p<0.05, **p<0.01, ***p<0.001 and ****p<0.0001; ^#^p<0.05, ^##^p<0.01, ^###^p<0.001 and ^####^p<0.0001; ^^ p<0.01.

## KEY RESOURCES TABLE

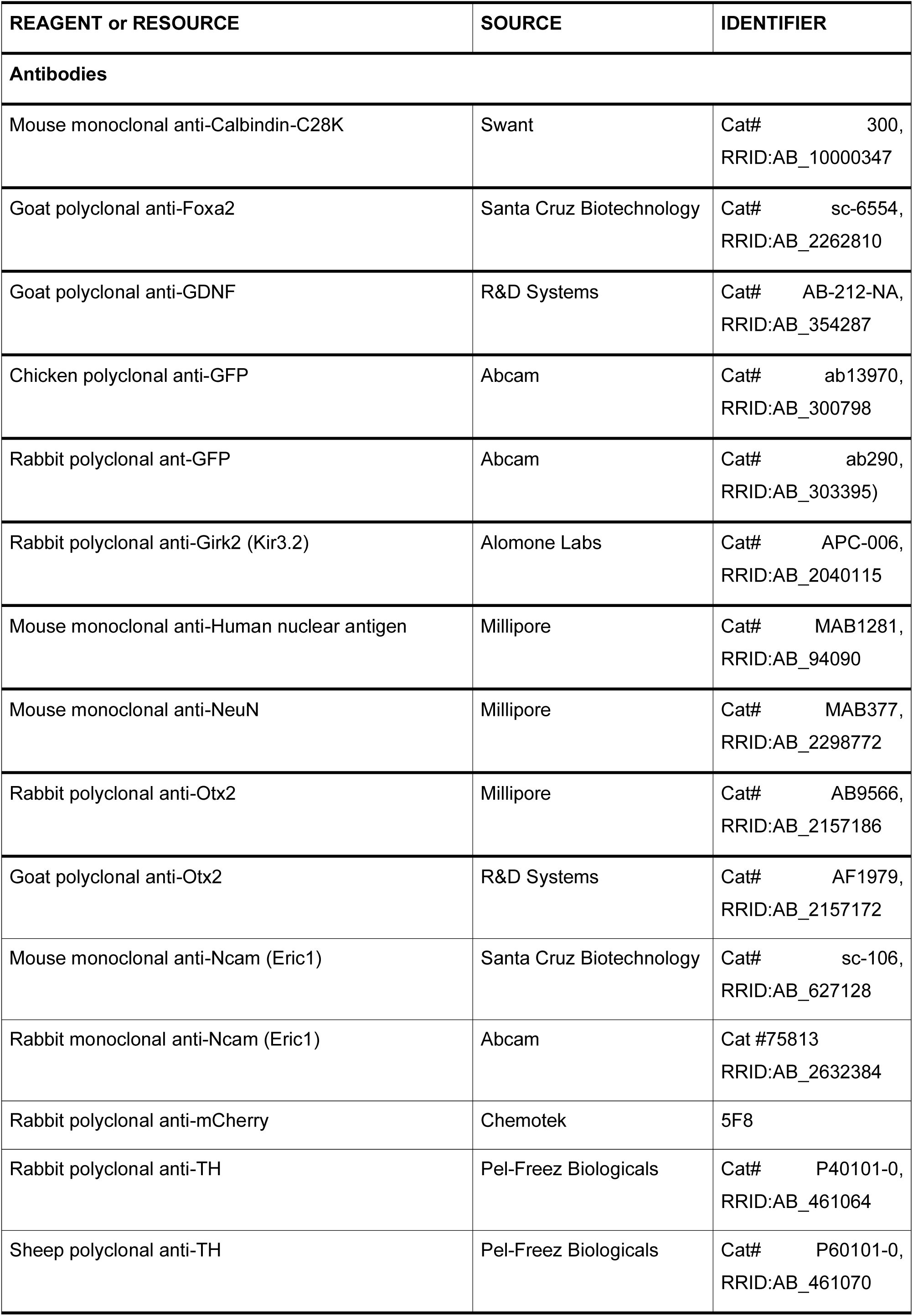
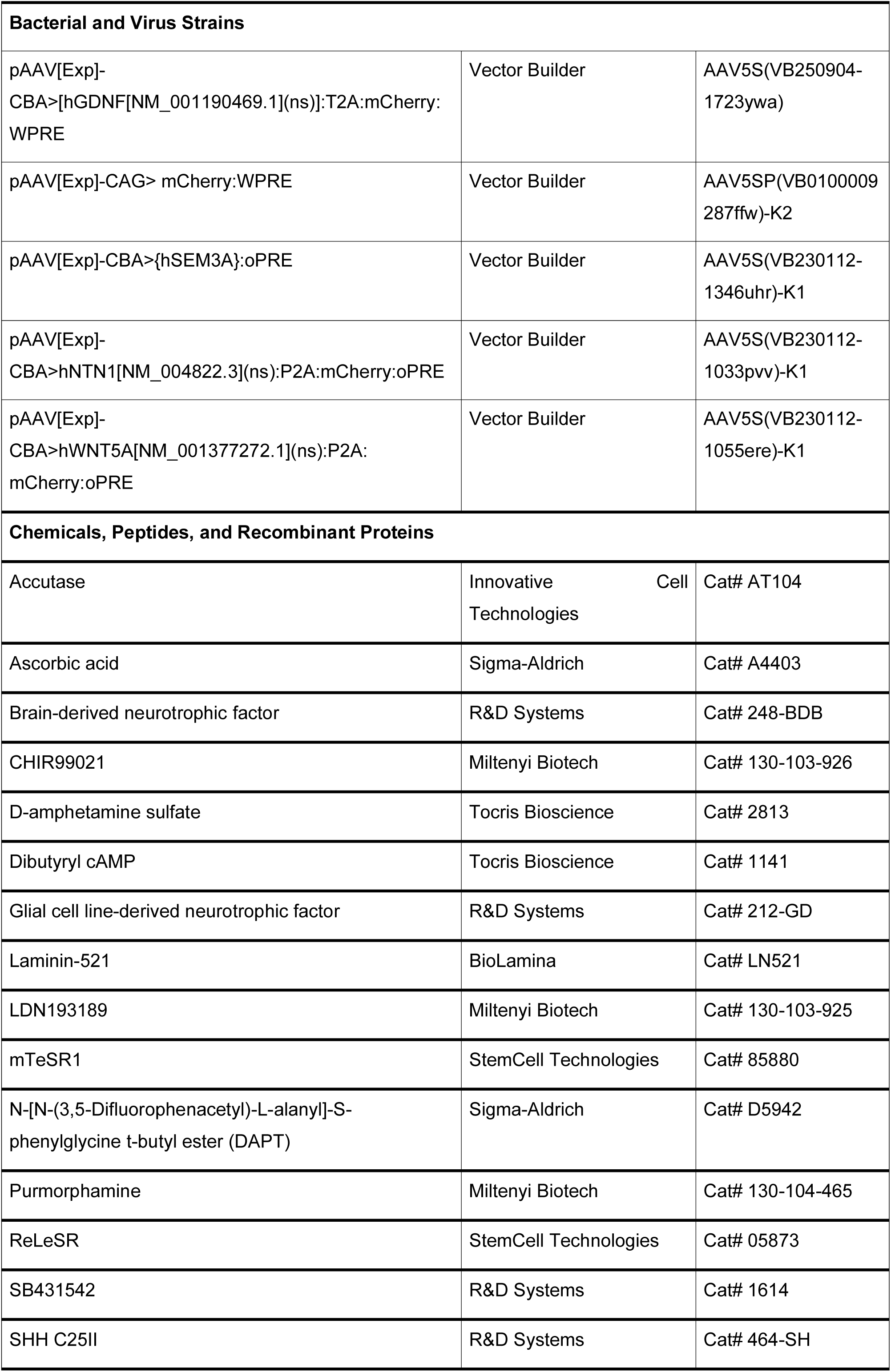
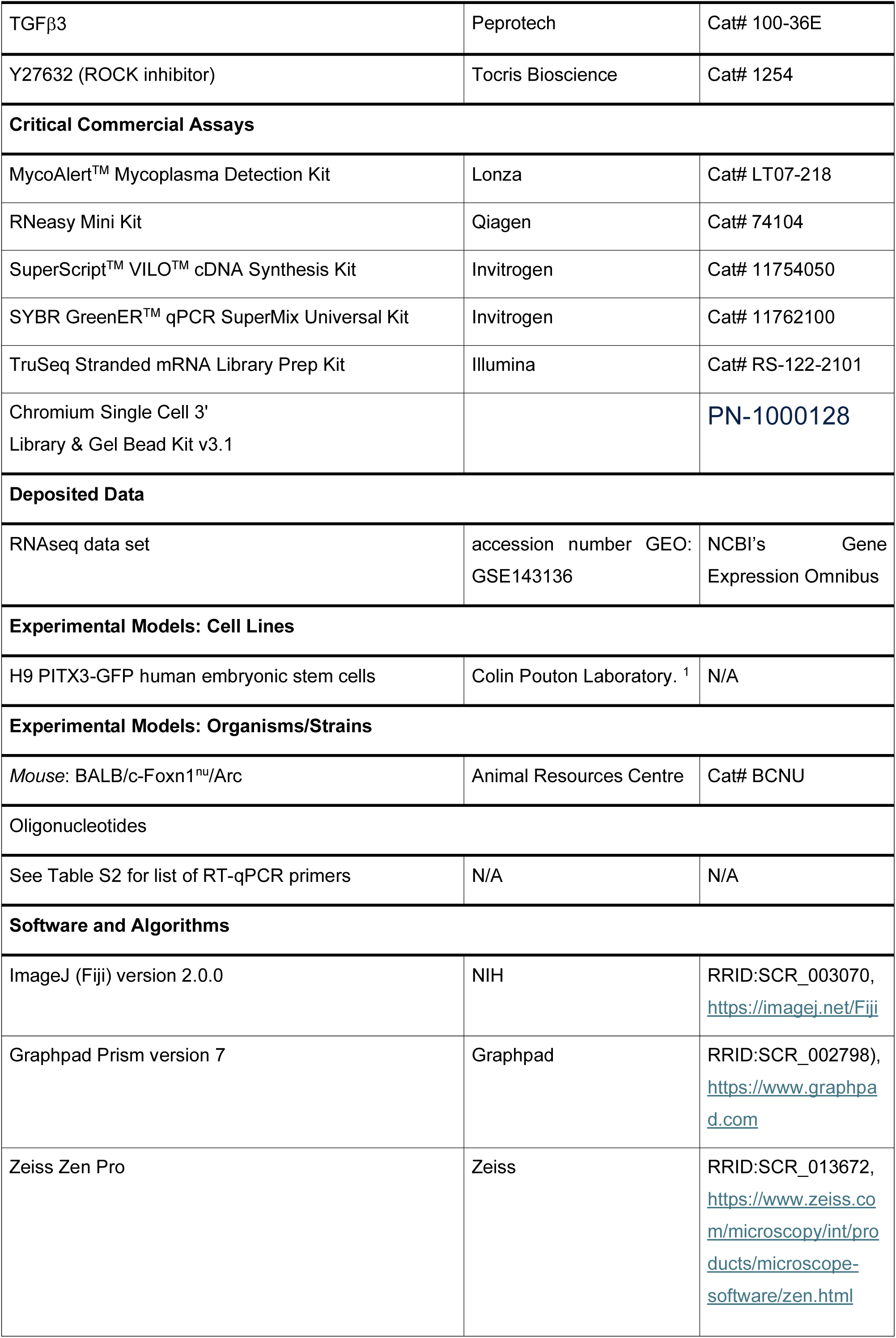
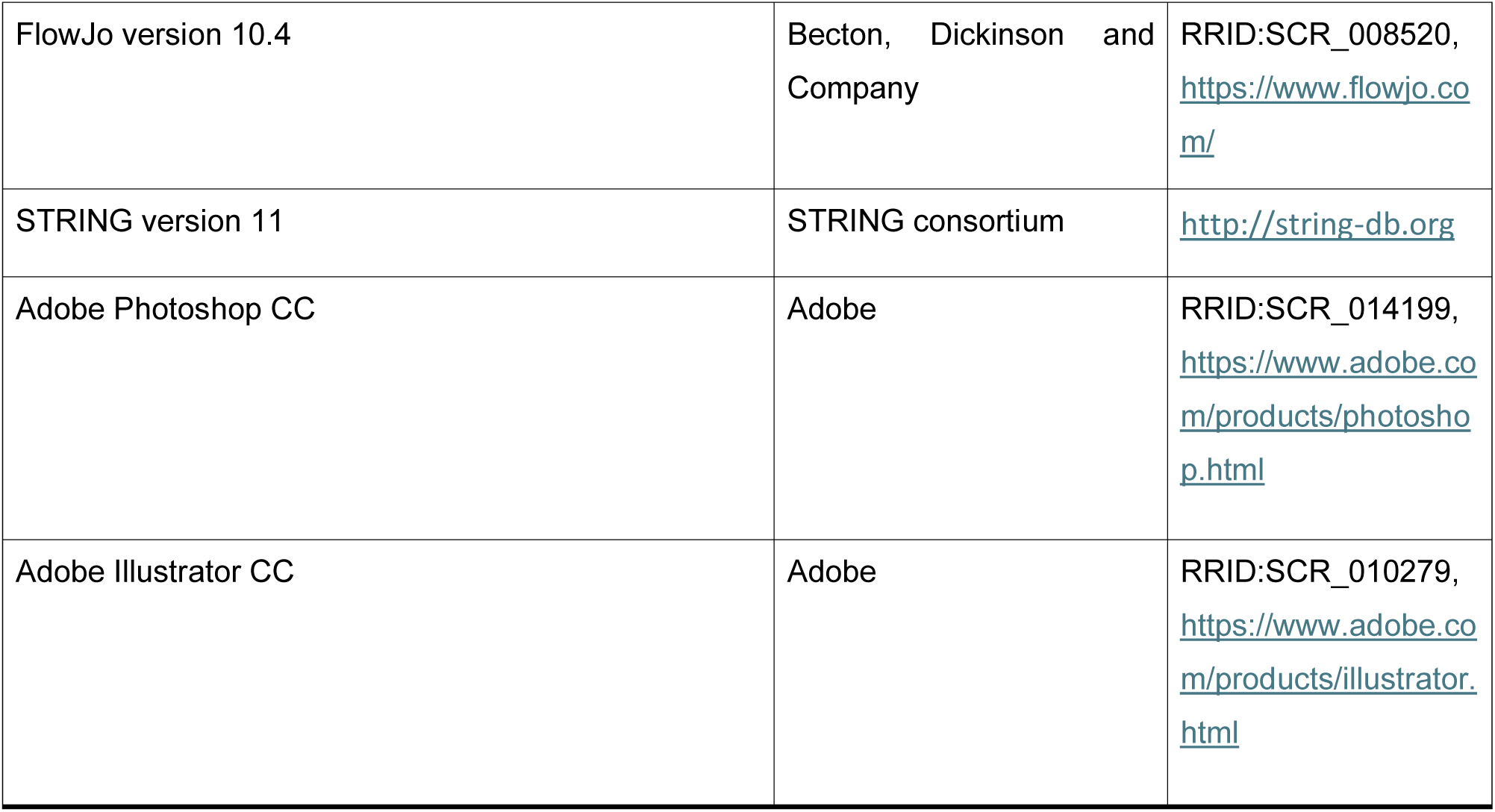

