## Supplementary Information for "Netrin-1 drives cell-type-specific plasticity of human dopaminergic neurons during circuit integration in a Parkinsonian model"

**SUPPLEMENTARY DATA**

**Supplementary Figure 1 – Tissue isolation, gating strategy and validation thereof used to separate human neuronal nuclei for single nuclei RNA sequencing (snRNA-seq).**

(A) Schematic showing site of cells injection from which graft survival was confirmed with (C') hematoxylin and eosin (H&E) staining. (B) Representative flow cytometry plot of samples used as negative controls to set up the flow cytometry gates for nuclei isolation: human pluripotent stem cells ($HNA^{+}$, $NeuN^{-}$), mouse brain tissue (HNA-) and unstained sample ($HNA^{-}$ $NeuN^{-}$).(C) Representative flow cytometry plot of human neuronal nuclei isolation using human nuclei (HNA) and neuronal nuclei (NeuN) staining. (D) Representative bright and fluorescent microscopy image showing intact isolated nuclei, no evidence of blebbing or lysis, and absence of live cells, debris, doublets, or clumps. (F) UMAP clustering of 5545 single-nuclei from the two xenografted rats revealed a clear separation between rat and human nuclei; nuclei are coloured by the fraction of mRNA transcripts aligned to human transcriptome. Abbreviations: A, area; FSC, forward scatter; H, height; HNA, human nuclear antigen; iPSC, induced pluripotent stem cells; NeuN, human neuronal antigen; SSC, side scatter; Tx, transplantation; VM DA, ventral midbrain dopaminergic neurons; W, width. Scale bars: (A) 100 µm

**Supplementary Figure 2- Single-nuclei RNA sequencing Quality Control.**

(A) Distribution of mitochondrial RNA % (mtRNA) and (B) features within sample 1 and sample 2; only nuclei that had a low proportion of mtRNA (<0.1%) i.e. were not apoptotic or lysing cells and high number of detected genes (>2000) passed the QC threshold and were included in the study. Dotted lines indicate thresholds for retaining high quality nuclei. (C) Proportion of human aligned transcripts; only nuclei in which >90% of mRNA were aligned to human transcripts were denoted as human origin. Dotted lines indicate thresholds for retaining human nuclei. (D) UMAP clustering of the human nuclei from the two xenografts showed no batch effect, as both samples identified two similar clusters. UMAP clustering of all the human nuclei showed no difference in (E) mtRNA % and (F) counts (number of genes/nuclei) between the clusters, meaning that the nuclei did not cluster based on mtRNA or number of identified genes. Abbreviations: mt, mitochondria. (G) ^^ score representing the most probable cell type in a cell cluster, showed that no clear cell-type could be ascribed to cluster 0. Dotted line indicates threshold for cell assignment. (H) Barplots of enriched subcellular localization of upregulated DEG's within grafted DA neurons; several upregulated genes were predicted to be a part of the synapse compartment. (I) Barplots of enriched molecular function and cellular component of upregulated DEG's within human DA neurons; upregulated genes were related to dopamine neuron and synapse functions e. g. dopamine binding, synapse part.

**Supplementary Figure 3 - Efficacy of the viruses to drive over-expression of proteins in vivo.**

(A-D) Mouse brain sections illustrating sustained mCherry (RFP+, A-C) or GDNF (D) expression in animals injected with (B) AAV-Control, (C) AAV-NTN1, (D) WNT5A or GDNF - indicative of protein expression within the host brain at 24 weeks after striatal injection of AAV (n=3 brains/group). (E) qPCR gene expression level of SEMA3A, NTN1 and WNT5A in homogenized mouse brain tissue 1 week after injection with AAV-SEMA3A, AAV-NTN1 or AAV-WNT5A, compared to the control (uninjected intact brain), n=4 brains/group. Data expressed as mean

±SEM. Scale bar: (A-D) 1 mm. Abbreviations: GDNF, glial cell-derived neurotrophic factor; NTN1, netrin; SEMA3A, semaphorin3A.

**Supplementary Figure 4 – Fixed frozen single nuclei sequencing quality control and annotation.**

(A) For each sample, QC was performed and only high quality single-nuclei were retained that had < 10% mitochondrial counts, between 500 and 10000 features and more than 1000 RNA counts. (B) Human and rat signature across the three different conditions (Ctrl, GDNF and NTN1). (C) Cluster analysis of the human nuclei from the three merged samples identified 16 distinct populations, represented across the samples with similar nCount_RNA and number of nuclei for each sample. (D) UMAP showing the 5 different populations identified and the distribution of key markers used to identify these populations as neurons (SYP, SYN1, GAP43, MAPT), astrocytes (GFAP, AQP4), oligodendrocytes (OLIG1), neural stem cells (SOX2) or mix. (E) Nuclei identified as neurons were re-clustered and nCount_RNA and number of nuclei were similar across the three samples. (F) Based on the expression of key dopaminergic, neuronal and non-neuronal markers nuclei were classified as non-dopaminergic or dopaminergic neurons. Abbreviations: CTRL, control; GDNF, glial cell-derived neurotrophic factor; NTN1, netrin.

**Supplementary table 1: List of viruses used in this study**

| **Viral Construct** | **Supplier** | **Cat#** |
|---|---|---|
| pAAV[Exp]-CAG> mCherry:WPRE | Vector Builder | AAV5SP(VB0100009287ffw)-K2 |
| pAAV[Exp]-CBA>[hGDNF[NM_001190469.1](ns)]:T2A:mCherry:WPRE | Vector Builder | AAV5S(VB250904-1723ywa) |
| pAAV[Exp]-CBA>{hSEM3A}:oPRE | Vector Builder | AAV5S(VB230112-1346uhr)-K1 |
| pAAV[Exp]-CBA>hNTN1[NM_004822.3](ns):P2A:mCherry:oPRE | Vector Builder | AAV5S(VB230112-1033pvv)-K1 |
| pAAV[Exp]-CBA>hWNT5A[NM_001377272.1](ns):P2A:mCherry:oPRE | Vector Builder | AAV5S(VB230112-1055ere)-K1 |

**Supplementary table 2: List of primers used in this study**

| Gene | Direction | Sequence |
|---|---|---|
| hNTN1 | FWD | ACTGCGATTCCTACTGCAAGGC |
| | REV | TTGTCCGCCTTCAGGATGTGGA |
| hWNT5A | FWD | TACGAGAGTGCTCGCATCCTCA |
| | REV | TGTCTTCAGGCTACATGAGCCG |
| hSEMA3A | FWD | GGTGCCTTATCAAGGAAGAGTCC |
| | REV | TACATGGCTGGATGACTTCTTGC |
| hGAPDH | FWD | CATGAGAAGTATGACAACAGCCT |
| | REV | AGTCCTTCCACGATACCAAAGT |

**Supplementary table 3: List of antibodies used in this study**

| Target | Host | Supplier | Cat# | Dilution |
|---|---|---|---|---|
| Calbindin-C28K | Mouse | Swant | CB300 | 1:1000 |
| cFOS | Goat | Santa Cruz | Cat# sc-52 | 1:1000 |
| DAPI | - | Sigma Aldrich | D9542 | 1:5000 |
| FOXA2 | Mouse | Abcam | Ab60721 | 1:4000 |
| FOXA2 | Goat | Santa Cruz | sc-6554 | 1:200 |
| GDNF | Goat | R&D Systems | Ab-212-NA | 1:1000 |
| GFP | Chicken | Abcam | Ab13970 | 1:1000 |
| GFP | Rabbit | Abcam | Ab290 | 1:200 |
| GIRK2 | Goat | Abcam | ab65096 | 1:500 |
| Human nuclear antigen (HNA) | Mouse | Millipore | MAB1281 | 1:300 |
| HNA-PE | Mouse | Abcam | Ab215755 | 1:200 |
| Nestin | Mouse | Millipore | MAB353 | 1:1000 |
| NeuN | Rabbit | Abcam | Ab236869 | 1:1000 |
| NeuN | Rabbit | Abcam | Ab104225 | 1:200 |
| OTX2 | Rabbit | Proteintech | 13497-1-AP | 1:200 |
| OTX2 | Goat | R&D Systems | AF1979 | 1:1000 |
| PSA-NCAM (Eric1) | Mouse | SantaCruz Biotechnology | sc-106 | 1:500 |
| PSA-NCAM (Eric1) | Rabbit | Abcam | ab75813 | 1:1000 |

| | | | | |
|---|---|---|---|---|
| mCherry | Rat | Chemotek | 5F8 | 1:1000 |
| TH | Mouse | ThermoFisher | TA506549 | 1:500 |
| TH | Rabbit | Peel-freeze | P40101-0 | 1:1000 |

**SUPPLEMENTARY FIGURES**

**PAVAN_Supp Figure 1**

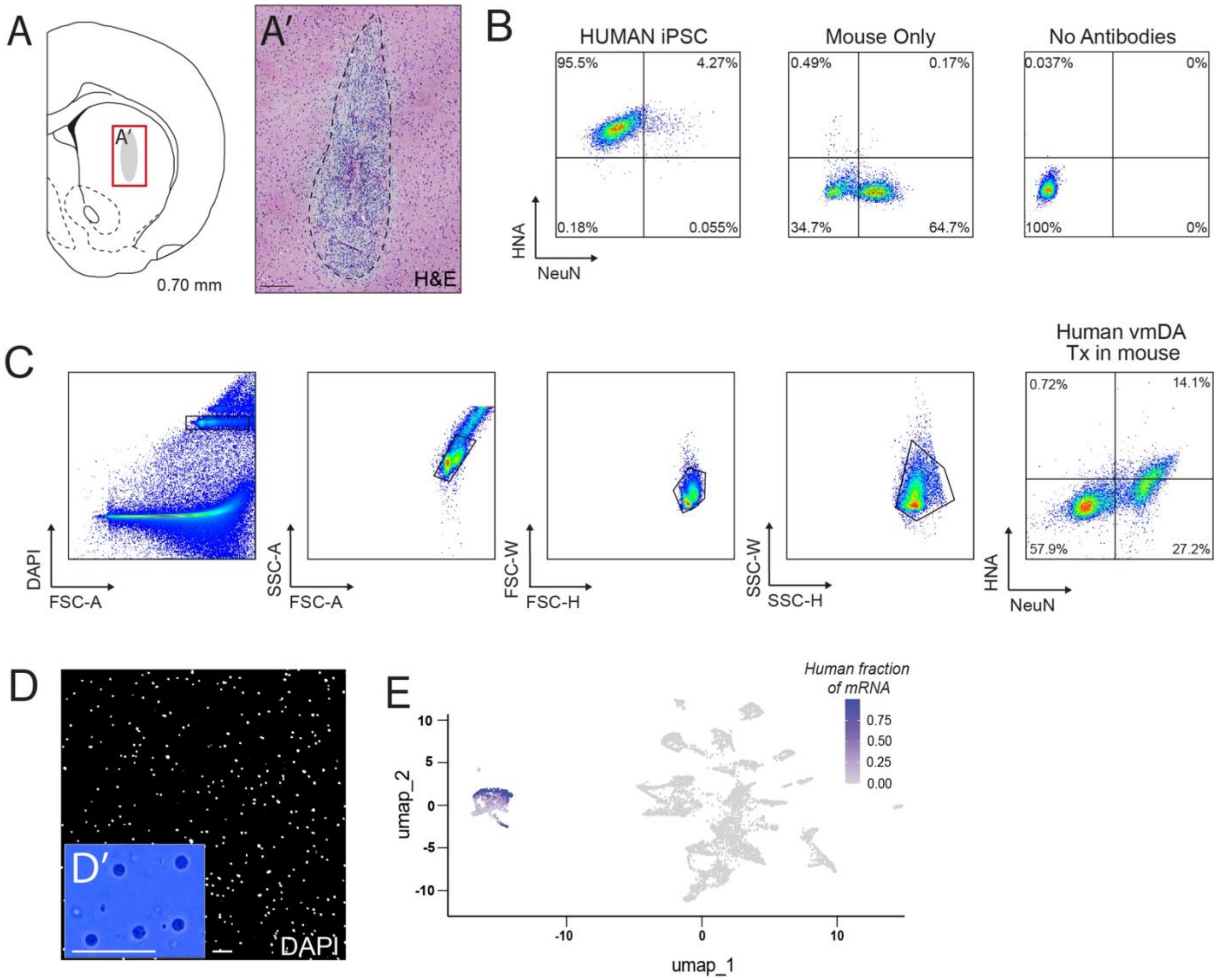

**PAVAN_Supp Figure 2**

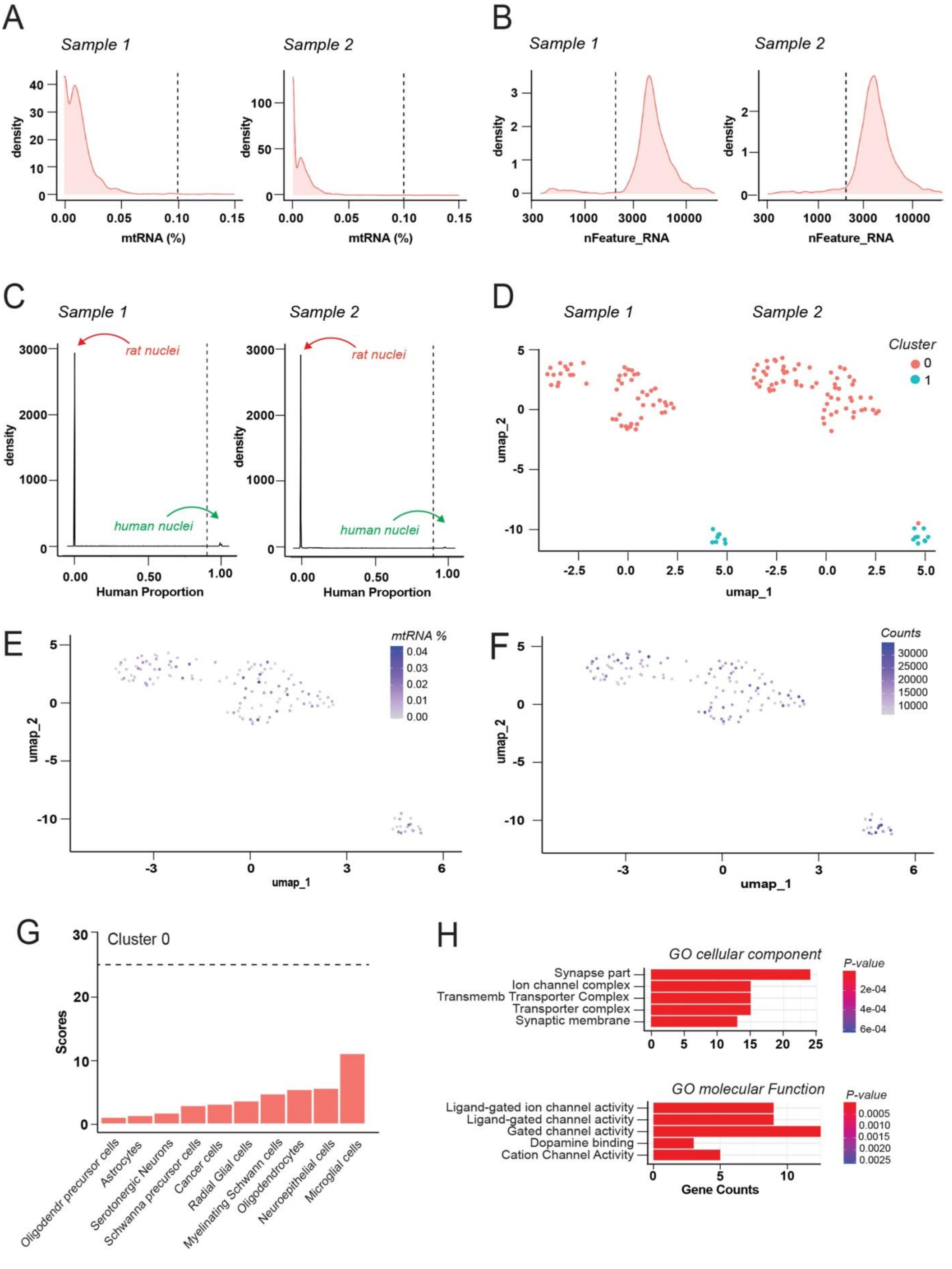

**PAVAN_Supp Figure 3**

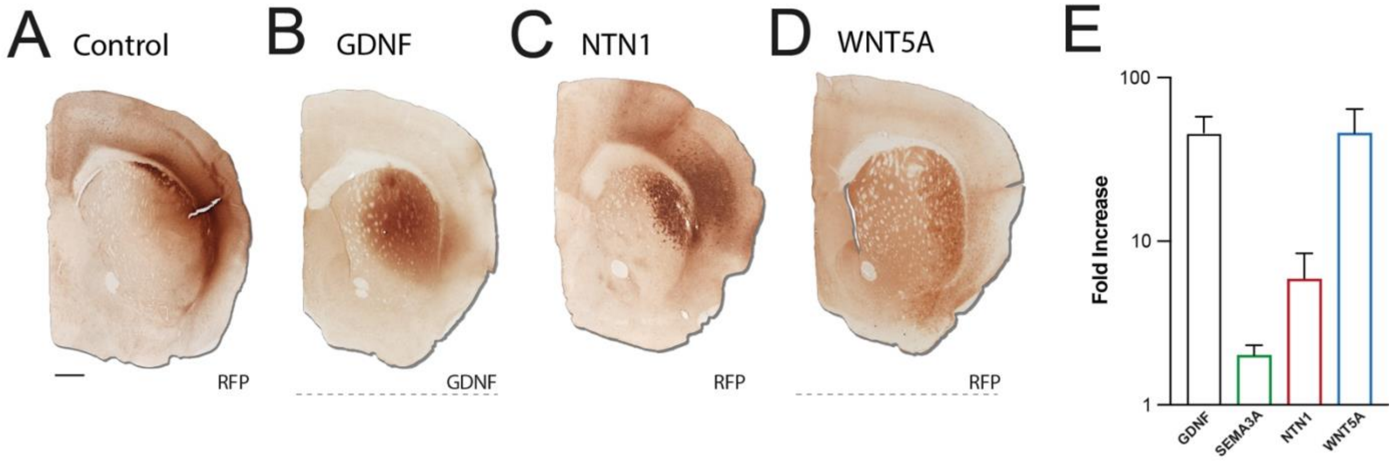

**PAVAN_Supp Figure 4**

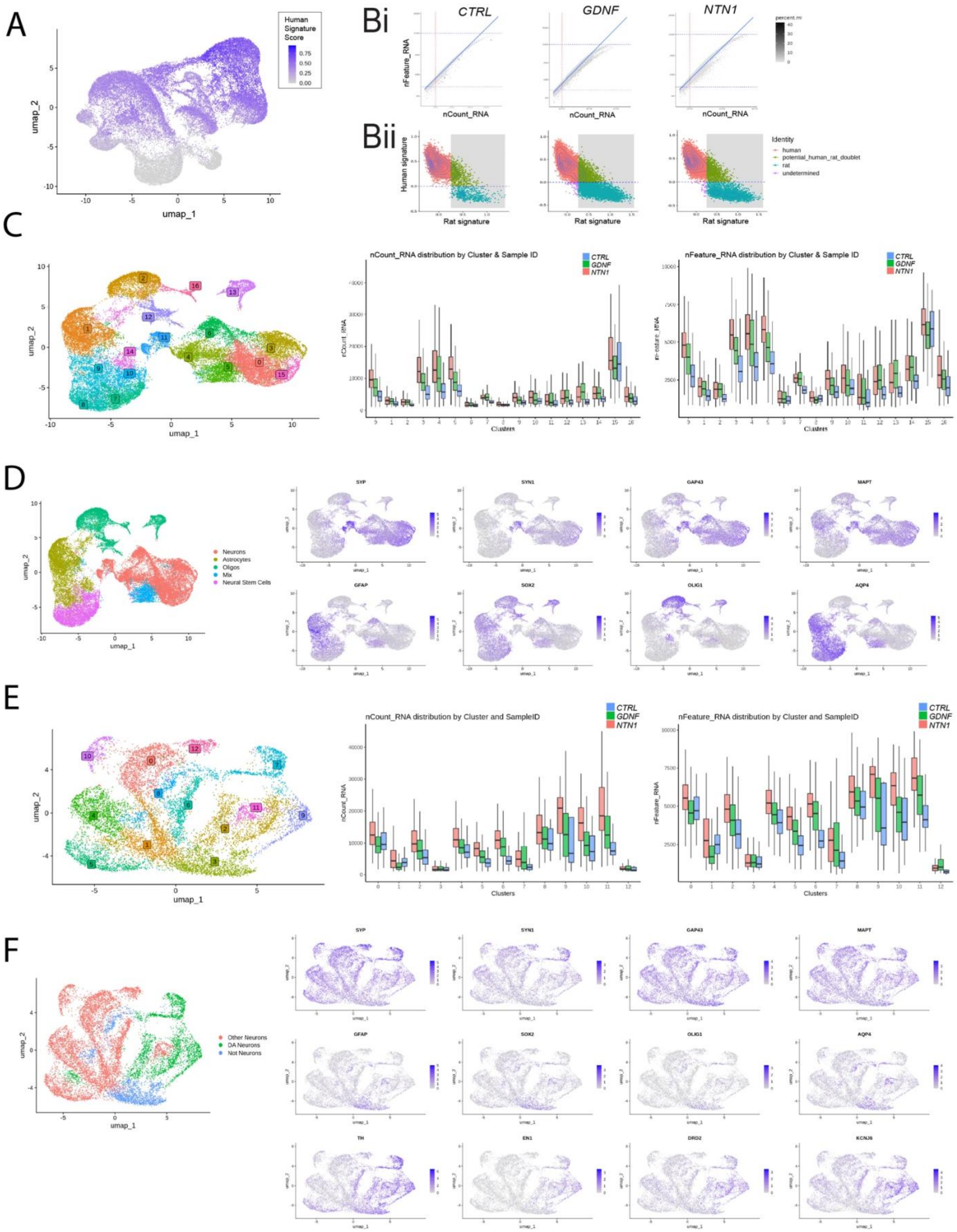
